# Accurate prediction of protein structures and interactions using a 3-track network

**DOI:** 10.1101/2021.06.14.448402

**Authors:** Minkyung Baek, Frank DiMaio, Ivan Anishchenko, Justas Dauparas, Sergey Ovchinnikov, Gyu Rie Lee, Jue Wang, Qian Cong, Lisa N. Kinch, R. Dustin Schaeffer, Claudia Millán, Hahnbeom Park, Carson Adams, Caleb R. Glassman, Andy DeGiovanni, Jose H. Pereira, Andria V. Rodrigues, Alberdina A. van Dijk, Ana C. Ebrecht, Diederik J. Opperman, Theo Sagmeister, Christoph Buhlheller, Tea Pavkov-Keller, Manoj K Rathinaswamy, Udit Dalwadi, Calvin K Yip, John E Burke, K. Christopher Garcia, Nick V. Grishin, Paul D. Adams, Randy J. Read, David Baker

**Affiliations:** Department of Biochemistry, University of Washington; Seattle, WA98195, USA; Institute for Protein Design, University of Washington; Seattle, WA98195, USA; Faculty of Arts and Sciences, Division of Science, Harvard University; Cambridge, MA02138, USA; John Harvard Distinguished Science Fellowship Program, Harvard University; Cambridge, MA 02138, USA; Eugene McDermott Center for Human Growth and Development, University of Texas Southwestern Medical Center; Dallas, TX, USA; Department of Biophysics, University of Texas Southwestern Medical Center; Dallas, TX, USA; Department of Biochemistry, University of Texas Southwestern Medical Center; Dallas, TX, USA; Howard Hughes Medical Institute, University of Texas Southwestern Medical Center; Dallas, TX, USA; Department of Haematology, Cambridge Institute for Medical Research, University of Cambridge; Cambridge, U.K.; Program in Immunology, Stanford University School of Medicine, Stanford, CA 94305, USA; Departments of Molecular and Cellular Physiology and Structural Biology, Stanford University School of Medicine, Stanford, CA 94305, USA; Molecular Biophysics & Integrated Bioimaging Division, Lawrence Berkeley National Laboratory, Berkeley, CA, USA; Department of Biochemistry, North-West University; 2531 Potchefstroom, South Africa; Department of Biotechnology, University of the Free State; 205 Nelson Mandela Drive, Bloemfontein, 9300, South Africa; Institute of Molecular Biosciences, University of Graz; Humboldtstrasse 50, 8010, Graz, Austria; BioTechMed-Graz; Graz, Austria; Department of Biochemistry and Microbiology, University of Victoria; Victoria, British Columbia, Canada; Life Sciences Institute, Department of Biochemistry and Molecular Biology, University of British Columbia; Vancouver, British Columbia, Canada; Howard Hughes Medical Institute, Stanford University School of Medicine, Stanford, CA 94305, USA; Howard Hughes Medical Institute, University of Washington; Seattle, WA98195, USA

## Abstract

DeepMind presented remarkably accurate protein structure predictions at the CASP14 conference. We explored network architectures incorporating related ideas and obtained the best performance with a 3-track network in which information at the 1D sequence level, the 2D distance map level, and the 3D coordinate level is successively transformed and integrated. The 3-track network produces structure predictions with accuracies approaching those of DeepMind in CASP14, enables rapid solution of challenging X-ray crystallography and cryo-EM structure modeling problems, and provides insights into the functions of proteins of currently unknown structure. The network also enables rapid generation of accurate models of protein-protein complexes from sequence information alone, short circuiting traditional approaches which require modeling of individual subunits followed by docking. We make the method available to the scientific community to speed biological research.

**One-Sentence Summary:** Accurate protein structure modeling enables rapid solution of structure determination problems and provides insights into biological function.

---

The combination of the outstanding performance of DeepMind at the CASP14 with the relatively little information provided on the method at the meeting prompted considerable questioning about if and when the method would become available to the scientific community, and whether such accuracy could be achieved outside of a world leading deep learning company like DeepMind. As described at CASP14 conference, the DeepMind AlphaFold2 methodological advances (1) included 1) starting from multiple sequence alignments (MSAs) rather than from more processed features such as inverse covariance matrices derived from MSAs, 2) replacement of 2D convolution with an attention mechanism that better represents interactions between residues distant along the sequence, 3) use of a two track network architecture in which information at the 1D sequence level and the 2D distance map level is iteratively transformed and passed back and forth, 4) use of an SE(3)-equivariant Transformer network to refine atomic coordinates generated from the two track network, and 5) end-to-end learning in which all network parameters are optimized by backpropagation from the final generated 3D coordinates through all the network layers back to the input sequence.

Intrigued by the DeepMind results, and with the goal of increasing protein structure prediction accuracy for structural biology research and advancing protein design (2), we explored network architectures incorporating different combinations of these five properties. Since no implementation details were provided, we experimented with a wide variety of approaches for passing information between different parts of the networks, as summarized in the Methods. We succeeded in producing a “two-track” network with information flowing in parallel along a 1D sequence alignment track and a 2D distance matrix track similar to that of AlphaFold2 in overall architecture, with considerably better performance than trRosetta (3) (BAKER-ROSETTASERVER and BAKER in Fig. 1B).

**Fig. 1.**
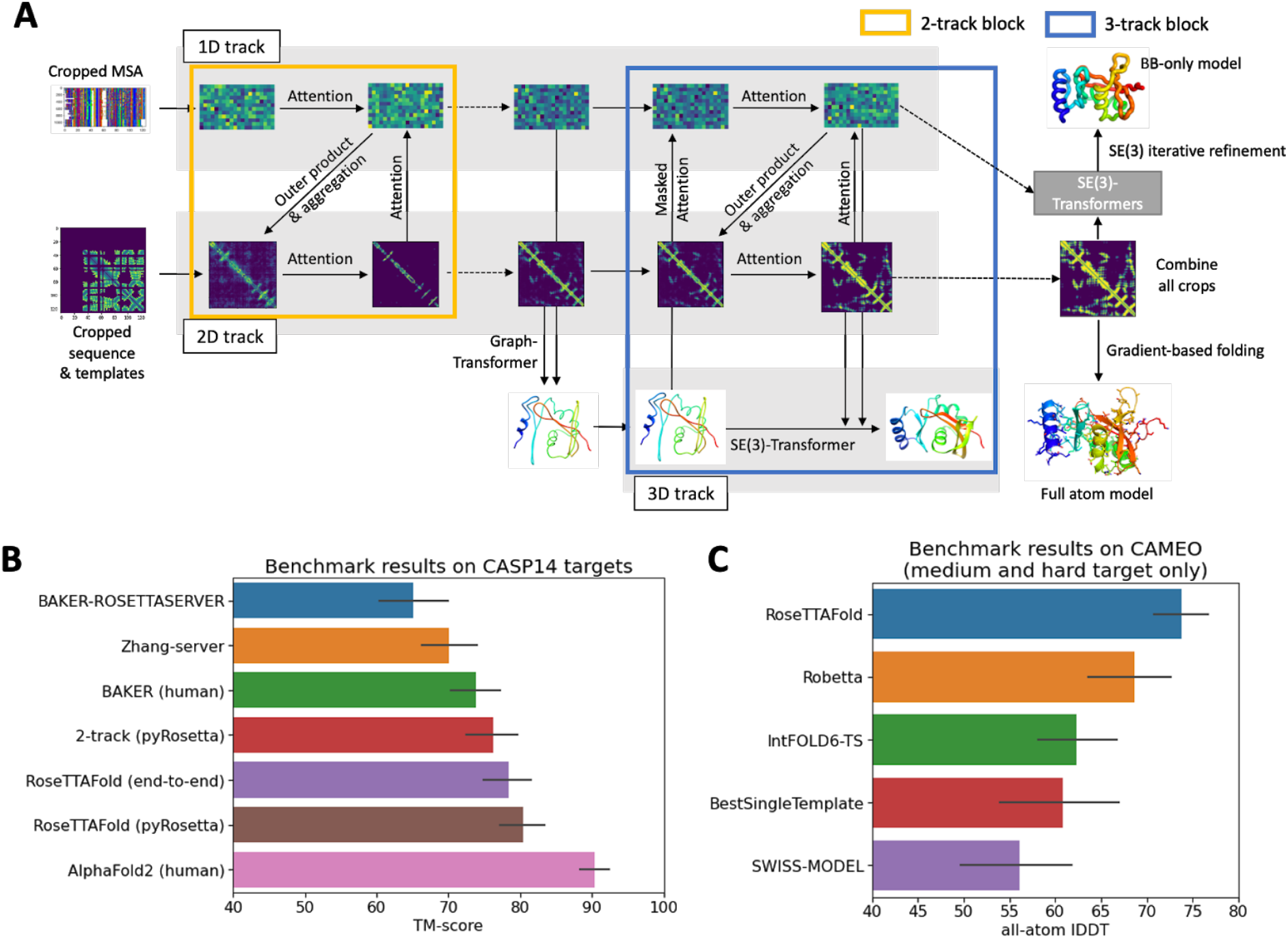
Network architecture and performance. (A) RoseTTAFold architecture with 1D, 2D, and 3D attention track. Each track frequently communicates to each other so that the network reason about relationships within and between sequences, distances, and coordinates simultaneously. See Methods and fig. S1 for details of each component. (B) Average TM-score (31) of all methods on the CASP14 targets. Zhang-server and BAKER-ROSETTASERVER were the top 2 server groups while AlphaFold2 and BAKER were the top 2 human groups in CASP14. Predictions of BAKER-ROSETTASERVER and BAKER were made based on trRosetta. Prediction with the 2-track model, and RoseTTAFold (both end-to-end and pyRosetta version) were completely automated. (C) Blind benchmark results on CAMEO. The model accuracy is evaluated based on all-atom lDDT (32) (data from CAMEO website: https://cameo3d.org/). For medium and hard targets having no clear template, RoseTTAFold outperformed the other servers.

We reasoned that still better performance could perhaps be achieved by extending to a third track operating in 3D coordinate space to provide a tighter connection between sequence, residue-residue distances and orientations, and atomic coordinates. We constructed architectures with the two levels of the two-track model augmented with a third parallel structure track operating on 3D backbone coordinates as depicted in Fig. 1A (see Methods and fig. S1 for details). In this architecture, information flows back and forth from the 1D amino acid sequence information, the 2D distance map, and the 3D coordinates, allowing the network to collectively reason about relationships within and between sequences, distances, and coordinates. In contrast, reasoning about 3D atomic coordinates in the AlphaFold2 architecture happens after processing of the 1D and 2D information is complete (although end-to-end training does link parameters to some extent). Because of computer hardware memory limitations, we could not train models on large proteins directly as the 3-track models have many parameters; instead, we presented to the network many discontinuous crops of the input sequence consisting of two discontinuous sequence segments comprising in total 260 residues. To generate final models, we combined and averaged the 1D features and 2D distance and orientation predictions produced for each of the crops and then used two approaches to generate final 3D structures. In the first, the predicted residue-residue distance and orientation distributions are fed into pyRosetta (4) to generate all atom models, and in the second, the averaged 1D and 2D features are fed into a final SE(3)-equivariant layer (5), and following end-to-end training from amino acid sequence to 3D coordinates, backbone coordinates are generated directly by the network (see Methods). We refer to these networks, which also generate per residue accuracy predictions, as RoseTTAFold. The first has the advantage of requiring lower memory (e.g. 8GB RTX2080) GPUs at inference time and producing full sidechain models, but it requires CPU time for the pyRosetta structure modeling step. For proteins over 400 residues, the end-to-end version requires 24G TITAN RTX GPU; we are currently extending this network to generate side chains as well.

The 3-track model with attention operating at the 1D, 2D, and 3D levels and information flowing between the three levels was the best model we tested (Fig. 1B), clearly outperforming the top 2 server groups (Zhang-server and BAKER-ROSETTASERVER), BAKER human group (ranked second among all groups), and our 2-track attention models on CASP14 targets. The performance of the 3-track model on the CASP14 targets was still not as good as AlphaFold2 (Fig. 1B). This could reflect hardware limitations that limited the size of the models we could explore, or more intensive use of the network for inference. DeepMind reported using several GPUs for days to make individual predictions, whereas our predictions are made in a single pass through the network in the same manner that would be used for a server; following sequence and template search (~1.5 hours) the end-to-end version of RoseTTAFold requires ~10 minutes on an RTX2080 GPU for proteins with less than 400 residues to generate backbone coordinates. The pyRosetta version takes 5 minutes for network calculations on a single RTX2080 GPU and an hour for full atom structure generation with 15 CPU cores. The relatively low compute cost makes it straightforward to incorporate the methods in a public server, and to predict structures for large sets of proteins, for example all human GPCRs, as described below.

We can draw a more definitive conclusion from the improved performance of the 3-track models over the 2-track model with identical training sets, similar attention-based architectures for the 1D and 2D tracks, and similar operation in inference (prediction) mode. This suggests that simultaneously reasoning at the multiple sequence alignment, distance map, and three-dimensional coordinate representations can more effectively extract sequence-structure relationships than reasoning over only MSA and distance map information.

Blind structure prediction tests are needed to assess any new protein structure prediction method, but CASP is held only once every two years. Fortunately, the CAMEO experiment (6) tests structure prediction methods blindly on protein structures as they are submitted to the PDB. RoseTTAFold has been evaluated since May 15th, 2021 on CAMEO; over the 28 medium and hard targets released during this time, it outperformed all other servers evaluated in the experiment (Fig. 1C).

We experimented with approaches for improving accuracy by more intensive use of the network during sampling. Since the network can take as input templates of known structure, we experimented with further coupling of 3D structural information and 1D sequence information by iteratively feeding the predicted structures back into the network as templates, reasoning that reasonably accurate 3D structural information could improve initial local structure prediction, among other properties. We also experimented with randomly subsampling from the multiple sequence alignments to sample a broader range of models. These approaches generated ensembles containing higher accuracy models, but the accuracy predictor was not able to consistently identify models better than those generated by the rapid single pass method (fig. S2). Nevertheless, we suspect that these approaches can systematically improve model accuracy and are carrying out further investigations along these lines.

With the recent considerable progress in protein structure prediction, a key question is what accurate protein structure models can be used for. We investigated the utility of the RoseTTAFold to facilitate experimental structure determination by X-ray crystallography and cryo electron microscopy, and for building models providing biological insights for key proteins of currently unknown structure.

Solution of X-ray structures by molecular replacement (MR) often requires quite accurate models. The much higher accuracy of the RoseTTAFold method than currently available methods prompted us to test whether it could help solve previously unsolved hard MR problems and improve the solution of borderline cases. Four recent crystallographic datasets that had eluded solution by MR using models available in the PDB were reanalyzed using RoseTTAFold models: glycine acyltransferase (GLYAT) from Bos taurus (fig. S3A), a bacterial oxidoreductase (fig. S3B), a bacterial surface layer protein (SLP) (Fig. 2A) and the secreted protein Lrbp from the fungus Phanerochaete chrysosporium (Fig. 2B). In all four cases the predicted models had sufficient structural similarity to the true structures that they led to successful MR solutions (see Methods for details; the per-residue error estimates allowed the better parts to be weighted more heavily).

**Fig. 2.**
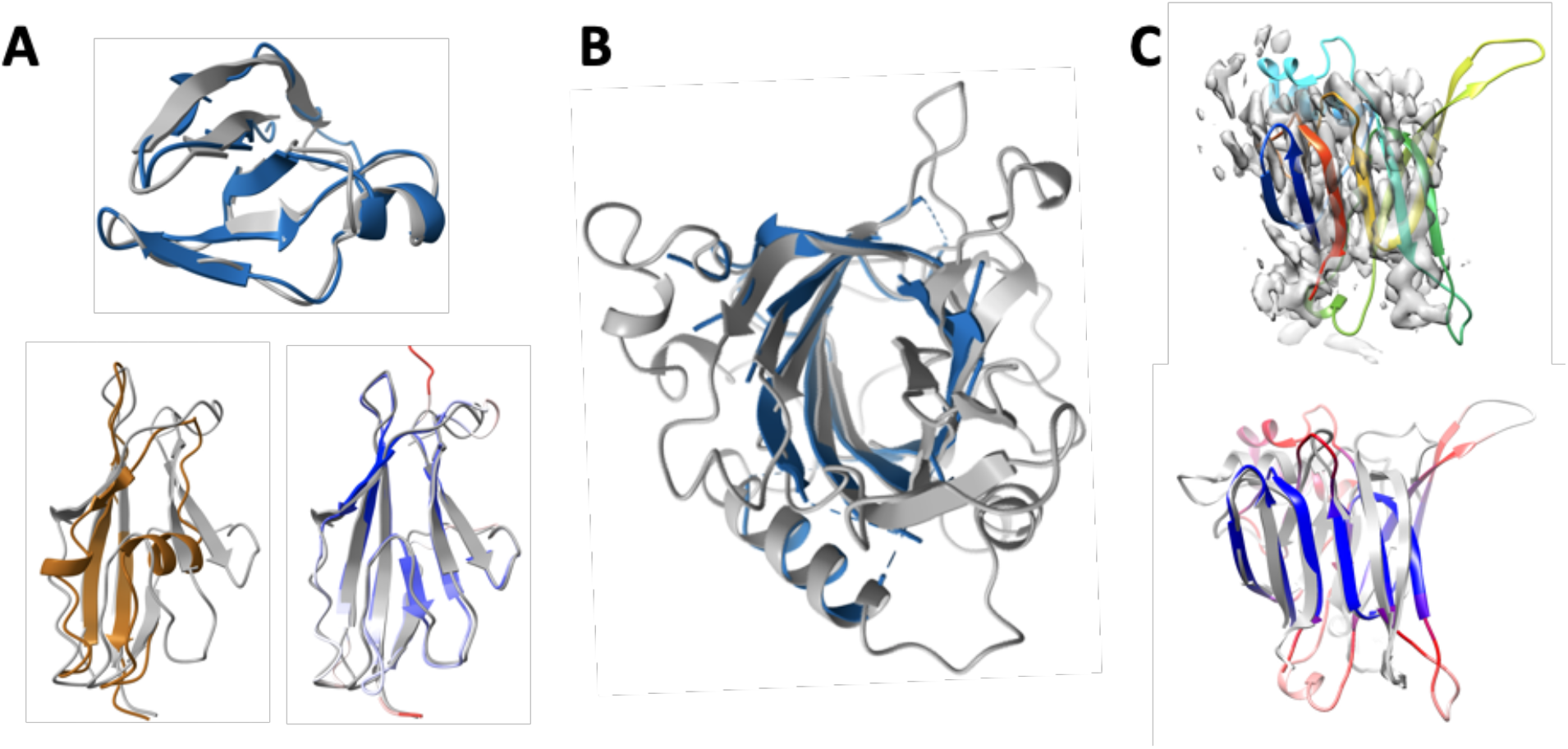
Enabling experimental structure determination with RoseTTAFold. **(A-B)** Successful molecular replacement with RoseTTAFold models. **(A)** C-terminal (top) and N-terminal (bottom) domain structure of the SLP. The final structure is shown in gray, with the only template for the N-terminal domain (PDB entry 4l3a) shown in brown. The RoseTTAFold model is colored blue for the C-terminal domain. For the N-terminal domain, the RoseTTAFold model is colored from blue to red by estimated RMS error, ranging from the minimum of 0.67 Å (blue) to 2 Å (red). **(B)** Refined structure of Lrbp (gray) with the best RoseTTAFold model superimposed. The RoseTTAFold model is colored in blue, and the residues having estimated RMS error greater than 1.3 Å are omitted. The full-length model of Lrbp is shown in fig. S3C. Cryo-EM structure determination of p101 G_βγ_ binding domain (GBD) in a heterodimeric PI3K_γ_ complex using RoseTTAFold. RoseTTAFold models colored in rainbow predicted a consistent all-beta topology with a clear correspondence to the density map (top). A comparison of the predicted and final structures is shown on the bottom. The RoseTTAFold model is colored from blue to red by estimated RMS error, ranging from the minimum of 1.50 Å (blue) to 3 Å (red). Figure prepared with ChimeraX (33).

To better understand why the RoseTTAFold models were successful, where PDB structures had previously failed, we compared the models to the crystal structures we obtained. The images in Fig. 2A and fig. S3 show that the homologue of the known structure closest to each of the unsolved targets was a much poorer model than the RoseTTAFold model of that protein; in the case of SLP, only a distant model covering part of the N-terminal domain (38% of the sequence) was available in the PDB, while no homologues of the C-terminal domain of SLP or any portion of Lrbp could be detected using HHsearch (7).

Building atomic models of protein assemblies from cryo-EM maps can be challenging if several of the subunits do not have homologues of known structure. We used RoseTTAFold to model the p101 G_βγ_ binding domain (GBD) in a heterodimeric PI3K_γ_ complex. The top HHsearch hit has a statistically insignificant E-value of 40 and only covers 14 residues out of 167 residues. The model could readily fit into the electron density map despite the low local resolution (Fig. 2C, top). trRosetta failed to predict a correct fold with the same MSA input (fig. S4). The RMSD between the structure prediction and final refined structure is 3.06 Å over the beta sheets (Fig. 2D, bottom; see Methods).

Experimental structure determination can provide considerable insight into biological function and mechanism. We investigated whether structures generated by RoseTTAFold could similarly provide new insights into function. We focused on two sets of proteins: first, G protein-coupled receptors of currently unknown structure, and second, a set of human proteins implicated in disease. Benchmark tests on GPCR sequences with determined structures showed that RoseTTAFold could generate accurate structures for both active and inactive states even in the absence of close homologs with known structures (fig. S5C; these models are considerably better than those in current GPCR model databases (8, 9)), and that the DeepAccNet model quality predictor (10) provided a good indication of actual model accuracy (fig. S5D). We generated models of closed and open states of all human GPCRs of currently unknown structure using RoseTTAFold, and we provide these along with their predicted per-residue accuracy.

Protein structures can provide insight into how mutations in key proteins lead to human disease. We identified human proteins with unknown structures that also lack close homologs of known structure and contain multiple disease causing mutations or have been the subject of intensive experimental investigation (see Methods). We used RoseTTAFold to generate models for 693 domains from such proteins. Over one-third of these models have an estimated lDDT > 0.8, which corresponded to an average RMSD of 1.5 Å on our known structure test set (fig. S5D). Here, we focus on three examples that illustrate the different ways in which structure models can provide insight into function or mechanisms of diseases.

Deficiencies in TANGO2 (transport and Golgi organization protein 2) lead to metabolic disorder, and the protein plays an unknown role in Golgi membrane redistribution into the ER (11, 12). The RoseTTAFold model of TANGO2 adopts an N-terminal nucleophile aminohydrolase (Ntn) fold (Fig. 3A) with well aligned active site residues that are conserved in TANGO2 orthologs (Fig. 3B). Ntn superfamily members with structures similar to the RoseTTAFold model suggest that TANGO2 functions as an enzyme that might hydrolyze a carbon-nitrogen bond in a bile acid related membrane component (13). Based on the model, known mutations which cause disease (magenta spheres in Fig 3A) could act by hindering catalysis (R26K, R32Q, and L50P, near active site) or produce steric clashes (G154R) (14) in the hydrophobic core. By comparison, a homology model based on very distant (<15% sequence identity) homologs had multiple alignment shifts that misplace key conserved residues (fig. S6)

**Fig. 3.**
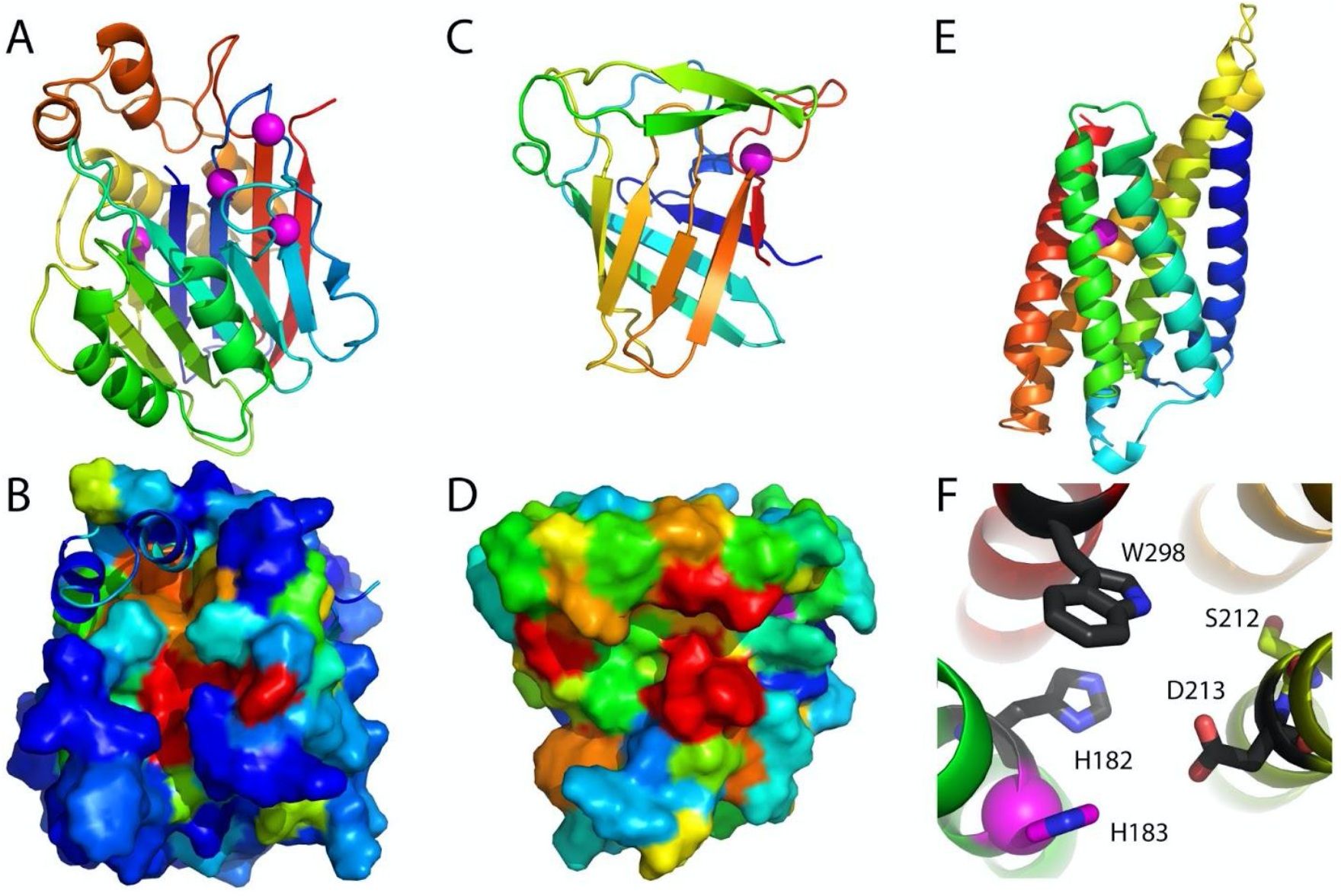
RoseTTAFold models provide insights into function. **(A)** TANGO2 model is colored in rainbow from the N-terminus (blue) to the C-terminus (red) and adopts an Ntn hydrolase fold with several pathogenic mutations (magenta spheres). **(B)** TANGO2 surface representation (~60° rotation about the x-axis with respect to panel (A) colored by ortholog conservation in rainbow scale from variable (blue) to conserved (red) highlights the active site. The surface is omitted for two helical extensions to the core Ntn fold that partially cover the active site. **(C)** ADAM33 prodomain adopts a lipocalin-like barrel shown in rainbow from N-terminus (blue) to C-terminus (red). The conserved position of a pathogenic mutation from ADAM10 is in the magenta sphere. ADAM33 model surface rendering is colored in rainbow by ortholog conservation from blue (variable) to red (conserved). The pathogenic mutation site is magenta. **(E)** CERS1 transmembrane structure prediction colored in rainbow from N-terminus (blue) to C-terminus (red), with a pathogenic mutation in TMH2 near a central cavity. **(F)** Residues that contribute to catalysis (H182 and D213) or are conserved (W298) line the cavity (black stick).

The ADAM (A Disintegrin And Metalloprotease) and ADAMTS families of metalloproteases are encoded by over 40 human genes, mediate cell-cell and cell-matrix interactions (15, 16) and are involved in a range of human diseases, including cancer metastasis, inflammatory disorders, neurological diseases and asthma (16, 17). The ADAMs contain prodomain and metalloprotease domains; the fold of the metalloprotease is known (18, 19), but not that of the prodomain, which has no homologs of known structure. The RoseTTAFold ADAM33 prodomain model has a lipocalin-like beta-barrel fold (Fig. 3C), whose extended superfamily includes metalloprotease inhibitors (MPIs) (20); this model is consistent with data suggesting that the ADAM prodomain inhibits metalloprotease activity using a cysteine switch (21); there is a cysteine in an extension following the predicted prodomain barrel. A pathogenic missense mutation (Q170H) in a related ADAM prodomain is near the cysteine switch (C173) (22); additional conserved residues within ADAM33 orthologs line one side of the barrel and likely interact with the metalloprotease (Fig. 3D), including another ADAM10 missense mutation (P139S) causing reticulate acropigmentation of Kitamura (23).

Transmembrane spanning Ceramide synthase (CERS1) is a key enzyme in sphingolipid metabolism using acyl-CoA to generate ceramides with various acyl chain lengths that regulate differentiation, proliferation, and apoptosis (24). Structure information is not available for any of the CerS enzymes, there are no homologs with known structures, and the number and orientation of its transmembrane helices (TMH) are not known (25). The RoseTTAFold CERS1 model for residues 98 to 304 (Pfam TLC domain) (26) includes six TMH that traverse the membrane in an up and down arrangement (Fig. 3E). A central crevice extends into the membrane and is lined with residues required for activity (His182 and Asp213) (27) or conserved (W298), as well as a pathogenic mutation (H183Q) found in progressive myoclonus epilepsy and dementia that decreases ceramide levels (28). This active site composition (His182, Asp 213, and potentially a neighboring Ser212) suggests testable reaction mechanisms for the enzyme (Fig. 3F).

The final layer of the end-to-end version of our 3-track network generates 3D structure models by combining features from discontinuous crops of the protein sequence (two segments of the protein with a chain break between them). Because the network can seamlessly handle chain breaks, it can be readily utilized to predict the structure of protein-protein complexes directly from sequence information. Rather than providing the network the sequence of a single protein, with or without possible template structures for this sequence, two or more sequences (and possible templates for these) can be input, with the output in this case the backbone coordinates of two or more disconnected protein chains. Thus, the network enables the direct building of structure models for protein-protein complexes from sequence information, short circuiting the standard procedure of building models for individual subunits and then carrying out rigid body docking. In addition to the great reduction in compute time required (complex models are generated from sequence information in ~30 min on 24G TITAN RTX GPU), this approach implements “flexible backbone” docking almost by construction as the structures of the chains are predicted in the context of each other. We tested the end-to-end 3-track network on paired sequences of complexes containing two chains and three chains from the PDB benchmark set (29) and found that the resulting models were very close to the actual structures as shown in Fig. 4 (panels A and B). Information on residue-residue co-evolution between the paired sequences likely contributes to the accuracy of the rigid body placement (fig. S7). The network was trained on monomeric proteins, not complexes, so there may be some training set bias in the structures of the monomers but there is none for the complexes. To illustrate the application of RoseTTAFold to complexes of unknown structure with more than three chains, we used it to generate models of the complete four chain human IL-12R/IL-12 complex (Fig. 4, panel C, fig. S8). A previously published cryoEM map of the IL-12 receptor complex indicated a similar topology to that of the IL-23 receptor, however, the resolution was not sufficient to observe the detailed interaction between IL-12Rβ2 and IL-12p35 (30). Such an understanding is important for dissecting the specific actions of IL-12 and IL-23 as well as generating inhibitors which block IL-12 without impacting IL-23 signaling. A computational model of the IL-12 receptor complex fits the experimental cryoEM density well and identified a shared interaction between Y189 in IL-12p35 and G115 in IL-12Rβ2 analogous to the packing between W156 in IL-23p19 with G116 in IL-23R. In addition, the model suggests a role for the IL-12Rβ2 N-terminal peptide (residue 24-31) in IL-12 binding not observed in the IL-12 cryo-electron microscopy (IL-12Rβ2 D26 may interact with nearby K190 and K194 in IL-12p35) and may provide an avenue to specifically target the interaction between IL-12 and IL-12Rβ2.

**Fig. 4.**
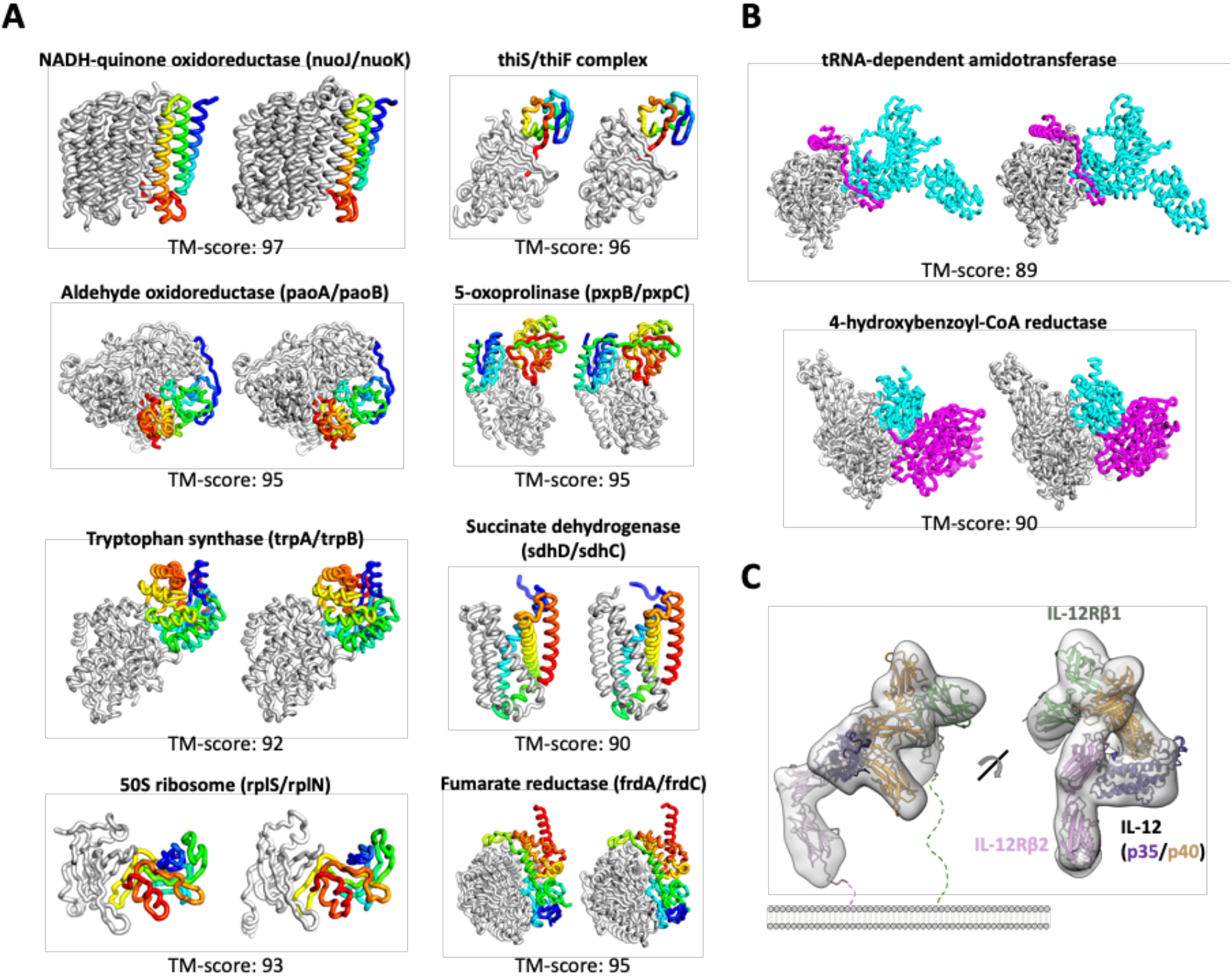
Complex structure prediction using RoseTTAFold. **(A)** Prediction of structures of *E*.*coli* protein complexes containing two chains from sequence information. The predicted structures (right) are close to the native structures (left). The first subunit is colored in gray and the second subunit is colored in rainbow from blue (N-terminal) to red (C-terminal). Complex TM-score is shown on the bottom. **(B)** Prediction of structures of protein complexes with three chains. RoseTTAFold predicted complex structures (right) are close to the native structures (left). Subunits are colored in gray, cyan, and magenta. **(C)** The IL-12R/IL-12 complex structures generated by RoseTTAFold. The RoseTTAFold model fits the density (EMD-21645) well.

RoseTTAFold enables solutions of challenging X-ray crystallography and cryo-EM modeling problems, provides insight into the functions of currently poorly understood proteins, and rapidly generates accurate models of protein-protein complexes. Further training on protein-protein complex datasets will likely further improve modeling of the structures of multiprotein assemblies. The approach can be readily coupled with small molecule and protein binder design methodology to improve computational discovery of new protein and small molecule ligands for targets of interest. The method is available as a server at https://robetta.bakerlab.org (RoseTTAFold option); access to rapid and accurate methods for prediction of protein structures and complexes should enable advances by biological researchers worldwide in many domains.

## Acknowledgments

We thank Eric Horvitz, Naozumi Hiranuma, David Juergens, Sanaa Mansoor, and Doug Tischer for helpful discussions, David E. Kim for web-server construction, and Luki Goldschmidt for computing resource management. TPK thanks Bernd Nidetzky and Mareike Monschein from Graz University of Technology for providing protein samples for crystallization. DJO gratefully acknowledges assistance with data collection from scientists of Diamond Light Source beamline I04 under proposal mx20303. TS, CB and TPK acknowledge the ESRF (ID30-3, Grenoble, France) and DESY (P11, PETRAIII, Hamburg, Germany) for provision of synchrotron-radiation facilities and support during data collection.

## Author contributions

MB, FD, and DB designed the research; MB, FD, IA, JD, SO, JW developed deep learning network; GRL and HP analyzed GPCR modeling results; QC, LNK, RDS, NVG analyzed modeling results for proteins related to the human diseases; CRG KCG analyzed modeling results for the IL-12R/IL-12 complex; PDA, RJR, CA, FD, CM worked on structure determination; AAD, ACE, DJO, TS, CB, TPK, MKR, UD, CKY, JEB, AD, JHP, AVR provided experimental data; MB, FD, GRL, QC, LNK, HP, CRG, PDA, RJR, DB wrote the manuscript; all authors discussed the results and commented on the manuscript.

## Data and materials availability

The GPCR models of unknown structures have been deposited to http://files.ipd.uw.edu/pub/RoseTTAFold/all_human_GPCR_unknown_models.tar.gz and http://files.ipd.uw.edu/pub/RoseTTAFold/GPCR_benchmark_one_state_unknown_models.tar.gz. The model structures for structurally uncharacterized human proteins have been deposited to http://files.ipd.uw.edu/pub/RoseTTAFold/human_prot.tar.gz. The atomic models have been deposited at the Protein Data Bank (PDB) with accession codes PDB: 7MEZ (full PI3K complex structure). The structures will be deposited in the PDB when completed.

## Supplementary Materials

Materials and Methods

Figs. S1 to S14

Tables S1 to S3

