## Supplementary material for "Accurate prediction of protein structures and interactions using a 3-track network": fig. S1

### Materials and Methods

#### Details of deep learning model

*Initial Embedding*

We describe the input multiple sequence alignments (MSA) as a matrix $x \in \mathbb{R}^{N\times L}$, where rows correspond to N sequences in the MSA, and columns are L positions in the aligned sequence. The input MSA is first tokenized to get the initial MSA features for further processing. Individual amino acids and gaps are regarded as character-level tokens (21 in total), and those are mapped to vectors having $d_{msa}$ size through an embedding layer. To let the network know the positional relationship between residues in each sequence, the sinusoidal positional encoding (34) is added for residues in each sequence. As an MSA is an unordered set of sequences, an indicator for the query sequence is added for the sequences instead of positional encoding in sequence dimension.

Template information is used to generate initial pair features by extracting pairwise distances and orientations from template structures for the aligned positions, along with 1D (positional similarity and alignment confidence scores) and scalar features (HHsearch probability, sequence similarity, and sequence identity) provided by HHsearch (7). Both features are concatenated to 2D inputs by tiling them along both axes of 2D inputs. Templates are first processed independently by one round of axial attention (row-wise attention followed by column-wise attention) (35) and then merged together into a single 2D feature matrix using a pixel-wise attention mechanism. This processed feature matrix is then concatenated with the 2D-tiled query sequence embedding and projected to hidden dimension ($d_{pair}$) for pair features. The 2D sinusoidal positional encoding is also added.

*Processing MSA features via self-attention*

After embedding the input MSA as described in the previous section, each MSA update step has $\mathbb{R}^{N\times L\times d}$ features as input and output. The MSA features are processed by the axial attention approach (35) which alternates attention over rows and columns of the 2D features. To reduce memory usage, we used Performer architecture (36) for the column attention (attention over sequence dimension) that reduces the memory requirements from $O(LN^2)$ to $O(LN)$. We first compared this MSA encoder with coevolution extractor (described in the next section) to the architecture with hand-crafted features (sequence profiles and inverse covariance matrices). As shown in Table S1 (architecture 1 vs 2), we found that having a learnable MSA encoder slightly improves distance and orientation prediction ($\Delta$loss=-0.07) as well as top L long-range contact accuracy by 2%p.

For the row attention (attention over residue dimension), we tested two different attention methods: 1) un-tied attention and 2) softly tied attention inspired by MSA Transformer architecture (37). In MSA Transformer, the tied attention idea for residue-wise attention was first introduced based on the fact that the homologous sequences in the MSA should have similar structures. Here, we modified this tied attention idea to reduce contributions from unaligned regions by introducing a learned position-wise weight factor (see Algorithm 1) to combine attention signals from sequences in MSA. We defined soft-tied attention to be Eq. (1) where N is the number of sequences in MSA, $Q_n$, $K_n$ are the matrix of queries and keys for n-th sequence of input, $W_n$ is the position-wise weight factor for the corresponding sequence.

$$attention = softmax(\sum_{n=1}^{N} W_n Q_n K_n^T) \quad \text{Eq. (1)}$$

In our experiments with small 2-track models, this soft-tied attention improves the top L long-range contact prediction performance by 2%p compared to the un-tied version (Table S1, architecture 6 vs 7). Interestingly, the soft-tied residue-wise attention maps showed correlations to the true contact map as shown in Fig. S9 (panel A and B). The final architecture used in RoseTTAFold is illustrated in Fig. S1A.

| Algorithm 1. Position-wise weight factor calculation |
| --- |
| Input:<br>● Q: embedding of query sequence (batch,1,L,$d_{msa}$)<br>● M: MSA embeddings (batch, N, L, $d_{msa}$)<br>● H: the number of attention heads for subsequent tasks<br><br>Get a query and key from given embeddings<br>Query = Linear($d_{msa}$, $d_{msa}$)(Q)<br>Key = Linear($d_{msa}$, $d_{msa}$)(M)<br><br>Permute & reshape Query and Key to calculate cross attention over sequence dimension<br>Query = permute_and_reshape(Query) # (batch, L, H, 1, $d_{msa}$//H)<br>Key = permute_and_reshape(Key) # (batch, L, H, N, $d_{msa}$//H)<br><br>Calculate attention between Query and Key<br>Attention = Query@Key.T # (batch, L, H, 1, N)<br><br>Take softmax for the last dimension<br>W = Softmax(Attention, dim=-1)<br><br>Output:<br>● W: positional weight for sequences |

*Update pair features with coevolution signal derived from MSA features*

To extract residue pairwise interaction information from given MSA features, we adopted the outer product and aggregation idea from the CopulaNet method (38). The outer product can capture the correlation between two residues in each sequence. By aggregating the signals from all sequences in MSA, we can measure the strength of covariation. For example, in the simplest case with one-hot encoded embedding for sequences, we get a 21x21 substitution table for each pair of positions (i,j) including gaps. When we take the average of the substitution tables from all sequences, the resulting 21x21 features will show different distributions depending on whether they interact with each other in 3D space or not. The broadly distributed 21x21 features indicate random uncorrelated mutations, and it means that those two residues are less likely to make contact in 3D space. On the other hand, if the aggregated features have sharp distributions (indicating correlated mutations), they will have a higher chance to directly interact with each other. In practice, the learned MSA embeddings through the network are used instead of one-hot encoding.

As outer products could require a huge memory ($O(d^2)$), the MSA embeddings are first projected down to the smaller hidden dimensions (32 features in this case) to reduce the memory requirements. After taking the outer product of embeddings derived from each sequence in MSA for any two residues, it calculates weighted averages of the outer products from all sequences with position-wise sequence weights. These aggregated coevolution features are then combined with 1D features (weighted average of MSA features) and residue-wise attention maps from the previous MSA update. They are projected down to match the hidden dimension for pair features.

To combine newly extracted pair features and previous pair features, we tested two different approaches: 1) Adding two pair features followed by feed-forward network and 2) concatenating two pair features followed by a single residual block of 2D convolutional network. As shown in Table S1 (architecture 5 vs 6), feature concatenation and 2D convolution clearly showed better performances, and we used this approach as outlined in Fig. S1B for our final model.

*Refine pair features via row and column-wise self-attention*

The updated pair features based on coevolution signals from MSAs are further refined by axial attention as shown in Fig. S1C. Using axial attention (35) instead of 2D convolution gave a clear improvement in inter-residue geometry predictions (Δloss=-0.35) with additional contact accuracy gain (Δaccuracy=2%p) even with single track architecture having sequential MSA and pair feature processing (Table S1, architecture 2 vs 3). This recapitulates one of DeepMind's observations that the attention mechanism is more suitable for protein structure prediction as it can directly learn the relationship between two residues distant in sequence.

In addition to the axial attention, we used Performer architecture (36) for attention to further reduce memory usage so that the larger architecture could fit on the GPU for experiments as larger architecture showed better performance (Table S1, architecture 7 vs 8).

*Update MSA features based on structure information encoded in pair features*

The most distinctive feature of AlphaFold2 architecture is that MSA features are updated based on the pairwise features. We experimented with two different ways to update MSAs based on given pair features: 1) taking cross-attention (or encoder-decoder attention) (34) between MSA and pair features by considering pair to MSA updates as a kind of encode-and-decode process 2) applying attention maps derived from pair features to MSA features (named direct-attention here) so that MSA features can be updated by attending positions close in 3D space that encoded in the pairwise features. As shown in Table S1 (architecture 4 vs 5), direct-attention showed clearly better performance (Δloss=-0.4, Δcontact accuracy=4%p). The attention maps derived from pairwise features showed a good agreement to the true contact map (Fig. S9, panel A and C). The final architecture based on direct-attention is outlined in Fig. S1D.

*Initial 3D structure prediction*

We employed Graph Transformer-based architecture (39) (shown in Fig. S1E) to generate initial backbone coordinates for 3D track (structure track). The input is defined as a fully connected graph with nodes representing the residues in the protein. The node and edge embeddings are learned from the averaged MSA features combined with one-hot encoded query sequence and the pair features along with sequence separation, respectively. The backbone coordinates are estimated using a stack of four Graph Transformer layers followed by simple linear transformation to predict Cartesian coordinates of N, $C_\alpha$, C atoms for each residue node.

*Structure updates through SE(3)-Transformer*

SE(3)-Transformer (5) is employed to refine the given xyz coordinate based on updated MSA and pair features in the 3-track model (Fig. S1F). The protein graph is defined with nodes representing $C_\alpha$ atoms, and each node is connected to the K-nearest neighbors. The positions of N and C atoms are encoded by including displacement vectors to the corresponding $C_\alpha$ atoms as the degree 1 node features (vector node features). The node embeddings derived from averaged MSA features and the one-hot encoded query sequence are used as degree 0 node features (scalar node features). Pair features corresponding to the graph edges are also included as input features for SE(3)-Transformer. SE(3)-Transformer predicts shifts of $C_\alpha$ atoms and new displacement vectors for N and C atoms to the updated $C_\alpha$ positions. It also gives degree 0 node features (called state features here) that are used to calculate attention maps for structure-based MSA updates described in the next section.

*Update MSA features based on a 3D structure*

Similar to the MSA updates based on pair features in the 2-track model, attention maps derived from the current 3D structures are used to update MSA features. Four attention maps are calculated based on the state features, and they are masked based on the $C_\alpha$ distances with four different cutoffs (8, 12, 16, and 20 Å) so that it only attends to the neighbors in 3D space. The same attention maps are applied to all the sequences in the MSA. The outputs from the masked multi-head attention are further processed by a pointwise feed forward layer. The entire process is outlined in Fig. S1G.

*Definition of 2-track and 3-track feature processing blocks*

We defined 2-track blocks with four arrows in Fig. 1A (orange box). It first updates MSA features through self-attention, extracts coevolution features from MSA and combines them with the previous pair features. Pair features are further optimized by axial attention, and MSA features are updated based on the structure information encoded in the current pair features. For 3-track blocks (blue box in Fig. 1A), we found that the order of communication between tracks is important. We experimented with two different ways to communicate 1D, 2D, and 3D tracks: updating structures before and after synchronizing MSA and pair features as shown in Fig. S10. The 3D coordinate updates based on synchronized MSA and pair features showed clearly better performance (Table S1, architecture 10 vs 11).

*Residue pairwise distance and orientation prediction*

Each of the inter-residue geometry representations (shown in Fig. S11) are predicted through a single residual block consisting of two 2D convolution layers with 3x3 filters followed by a convolution with 1x1 filters and softmax activation. Since maps for $C_\beta$-$C_\beta$ distances and dihedral angles along pseudo $C_\beta$-$C_\beta$ bonds are symmetric, we enforce symmetry in the network by using averages of transposed and untransposed feature maps as inputs for those predictions.

*Additional structure module for iterative refinement through the network*

Although structures are explicitly sampled in 3-track blocks, an additional structure module is introduced to build a model based on combined 1D features and 2D inter-residue geometry predictions for inference with multiple discontinuous crops. Initial coordinates for backbone N, $C_\alpha$, C atoms are generated using simple graph-based architecture (see Initial 3D structure prediction section above) with node and edge features derived from averaged MSA features and 2D distance and orientation distributions. These coordinates are further refined with multiple SE(3)-Transformer layers (5) by taking the same node and edge features used to generate initial coordinates. At the end of SE(3)-Transformer layers, the residue-wise $C_\alpha$-lDDT (32) is also estimated based on the degree 0 features from the final SE(3)-Transformer layer.

During the training, we didn't use any iteration, and the parameters were optimized through a single pass of the network. However, we found that we could use this structure module as an iterative refinement tool by feeding the output coordinates of the final SE(3)-Transformer layer to the first SE(3) layer as inputs at inference time (Fig. S12). The predicted $C_\alpha$-lDDT is used as a scoring function to decide when to stop the iteration and select the final model from all the sampled structures.

*Comparison between 2-track end-to-end model and 3-track model*

AlphaFold2 passed information from the 2-track trunk model into a 3D equivariant network operating on 3D coordinates directly. AlphaFold2 also employed end-to-end training, updating all parameters of the model by backpropagation from a loss function computed on 3D coordinates after many SE(3)-equivariant layers. As an experiment, we built a model with SE(3)-Transformer layers on top of the graph-based initial coordinate generation following the 2-track model. We found that adding SE(3)-Transformer layers improved the accuracy of structures generated by the simple graph-based network (Fig. S13), but this 2-track end-to-end model was not as good as the 3-track end-to-end model (Table S1, architecture 9 vs 12).

*Training details*

The extended trRosetta training set (containing 22,922 clusters with sequence identity cutoff 30%, 208,659 protein chains released in the PDB as of 02/17/2020) was used to train RoseTTAFold. Every training epoch, we cycled through all sequence clusters by picking a random protein chain from each cluster. For each selected protein chain, a subsampled MSA (having maximum NxL=214 tokens) and up to 10 randomly selected templates were used to augment training data. During training, protein chains over 260 residues in length were cropped to fit into GPU memory.

The loss function used to train the model consists of 1) distance and orientation prediction loss (cross entropy) with 0.5 Å and 10° bins (3), 2) coordinate and distance RMSD of predicted coordinates, and 3) mean squared error of predicted $C_\alpha$-lDDT score. During training, weights for coordinate and distance RMSD were ramped up from 0.05 to 0.2. For the other loss terms, weights are set to 1.0.

We train 130M parameters models having eight 2-track blocks and five 3-track blocks. Using eight 32GB V100 gpus, it took about 4 weeks to train the model upto 200 epochs. The following hyper-parameters were used:

- MSA, pair, template embedding size: 384, 288, and 64, respectively
- The number of attention heads for self-attention on MSA, pair, and template: 12, 8, and 4
- The number of attention heads for MSA updates based on pair features: 4
- Size of node and edge features for initial coordinate generation: 64
- The number of attention heads for initial coordinate generation: 4
- Size of input node and edge features for SE(3)-Transformer: 32
- SE(3)-Transformer architecture: 2 layers with 16 channels, 4 attention heads, and up to representation degree 1 (l=0 and 1 features were used)
- The number of closest residues to define graph for SE(3)-Transformer: 128 for first two 3-track blocks, 64 for last three 3-track blocks
- Learning rate: 0.0005 with linear learning rate decay after 16000 warmup steps
- Effective batch size: 64 in total (8 V100 GPUs, single training example per GPU, 8 gradient accumulation steps)
- Weight decay: 0.0001

**RoseTTAFold modeling pipeline**

We built a fully automated modeling pipeline based on RoseTTAFold. It first iteratively searches homologous sequences against UniRef30 (*40*) and BFD (*41*) sequence databases using HHblits (*7*). The E-value cutoff for sequence search is gradually relaxed until the resulting MSA has at least 2000 sequences with 75% coverage or 5000 sequences with 50% coverage (both at 90% sequence identity cutoff). The generated MSA is used to perform template searches against the PDB100 database with HHsearch (*7*).

With MSA and top 10 templates as input, the RoseTTAFold network predicts inter-residue geometries (probability distributions of 6D coordinates described in Fig. S11) for many 300×300 discontinuous crops (150 residues per each segment) and combined them by taking weighted averages based on predicted $C_\alpha$-lDDT values. We used two different strategy to generate final structure model with this combined 6D coordinate distribution: 1) gradient-based folding using pyRosetta (*4*) script and 2) structure module based on SE(3)-Transformer architecture described above (see *Additional structure module for iterative refinement through the network* section). The first method doesn't require large memory GPU as it predicts 300×300 size of 6D coordinates only, and gives full-atom model at the end, but it requires more CPU cores and time to run multiple trajectory (15 in total) of gradient-based folding from scratch. The second method can model backbone coordinates much faster than gradient-based folding (with similar accuracy level), but it requires a large memory GPUs (e.g. TITAN RTX) for proteins having more than 400 residues.

For the pyRosetta-based modeling protocol, the five models out of 15 sampled structures are selected based on predicted lDDT of DeepAccNet (*10*) after clustering. The $C_\alpha$ RMS error is estimated by converting predicted non-local $C_\alpha$-lDDT (only considering residue pairs having sequence separation > 12) using Eq. (2). This pyRosetta-based protocol is implemented in the Robetta server.

$$C_\alpha\ RMS\ error = 1.5e^{4\times(0.7-lDDT)} \quad \text{Eq. (2)}$$

### Molecular replacement calculations

*Structure of glycine N-acyltransferase*

The structure of glycine N-acyltransferase (GLYAT) from *Bos taurus* had evaded numerous attempts at solution, in spite of the availability of excellent data from three crystal forms. Structures of homologues were found using HHpred (*42*), which revealed that the only known structures were from distant relatives, almost all with low coverage of the target. Only 3 homologues (including the top hit) had greater than 60% coverage; these were only 12% identical in sequence. The top 5 hits were prepared for molecular replacement trials by pruning non-conserved side chains and loops using phenix.sculptor (*43*). In addition, an ensemble model was prepared by superimposing the individual homologues in phenix.ensembler (*44*) and trimming parts of the ensemble that are poorly conserved to leave a small conserved core. Molecular replacement trials with Phaser (*45*), MoRDa (*46*) and I-TASSER-MR (*47*) on all three crystal forms, using individual models, ensemble models and domain models, failed to yield any convincing results.

In contrast, molecular replacement was straightforward for all three crystal forms when using the RoseTTAFold models, whether as individual models or trimmed ensembles. An estimate of the effective RMS error is required to calibrate the likelihood target, and a value of 1.2 Å was used for these models.

A post mortem analysis was carried out to verify that model quality was the limiting factor for molecular replacement with models derived from the PDB. This analysis concentrated on a tetragonal crystal form, which diffracts to 1.5 Å resolution and has a single copy in the asymmetric unit. The other two crystal forms each have two copies of the protein in the asymmetric unit.

In the likelihood-based molecular replacement algorithm implemented in Phaser, the log-likelihood-gain (LLG) score is an excellent predictor of success. If LLG scores of 60 or more are achieved in placing a single copy, the solution is almost always correct (*48*). In contrast, scores below 30 are more likely to correspond to random incorrect placements. By correctly positioning a molecular replacement model and carrying out a rigid-body refinement in Phaser, we can evaluate the score that could have been achieved in the search. This calculation shows that none of the available models came close to providing sufficient signal to solve the structure, giving LLG scores of only 7 to 11 when correctly placed. The best model (with a score of 11) was the top hit in HHpred, PDB entry 1sqh. A full molecular replacement search with this model yielded a top LLG score of 22 for an incorrect placement. The high quality of the RoseTTAFold model, especially compared to the model derived from 1sqh, can be seen in Fig. S3A. For this figure, the experimental structure is illustrated using chain A from the current model of the hexagonal crystal form, in which the poorly ordered loop is most clearly defined. Table S2 summarizes the refinement statistics for this structure, as well as the oxidoreductase discussed below.

*Value added by coordinate error estimates for GLYAT structure determination*

LLG scores obtained with the RoseTTAFold models were compared, either ignoring the estimates provided for the RMS error of each amino acid or using it to weight each atom's contribution by providing a B-factor equal to $8\pi^{2/3}RMS^2$(*49*). Before applying the B-factor weighting, the LLG scores ranged from 88 to 148 for the 5 alternative models. After applying the weighting, the LLG scores ranged between 117 and 188. Similarly, the LLG score for the trimmed ensemble model increased from 191 to 244. In a more marginal case, such weighting could well be pivotal to success. Fig. S3A illustrates the correlation between predicted and actual errors in the RoseTTAFold, especially in the poorly ordered loop which has the highest predicted errors.

*Structure of a bacterial oxidoreductase*

The structure of an oxidoreductase from a bacterial source wasn't solved by molecular replacement using related structures available from the PDB, identified using HHpred (*42*). These efforts were likely unsuccessful because available structures had low sequence identity and only moderate sequence coverage - the best structures had an identity of ~33% for the first 40% of the sequence, or ~25% identity for the first 60% of the sequence. In addition, the 2 crystal forms were expected to have 6 or 12 molecules in the crystallographic asymmetric unit based on the most probable solvent content. The top 5 HHpred structures were prepared for molecular replacement trials by pruning non-conserved side chains and loops using phenix.sculptor (*43*). In addition, an ensemble model was prepared by superimposing the individual homologues in phenix.ensembler (*44*) and trimming parts of the ensemble that are poorly conserved to leave a small conserved core. Molecular replacement trials with Phaser (*45*) did not produce correct solutions as judged by significant overlaps between placed molecules, and a modest TFZ score of 7.4 in the lower probability P2 space group.

The top 5 RoseTTAFold models were superimposed using phenix.ensembler and parts of the ensemble that are poorly conserved were automatically trimmed. Atomic B-factors were calculated from the estimated RMS error as described above. Molecular replacement trials with Phaser produced a solution in the more likely $P2_1$ space group, albeit with a modest TFZ score of 6.9. Manual inspection of the solution revealed that 4 of the molecules formed 2 dimers, which were expected on the basis of the closest homologue structures and biophysical data. One dimer was extracted from the model and used in a new MR trial, which produced a very clear solution with a TFZ of 17.2. Comparison of the 2 molecular placement trials showed that the initial search had placed 5 molecules correctly but the 6th incorrectly. The successful dimer-based solution was used as the starting point for phase improvement using statistical density modification methods (*50*) in Phenix (*51*). The resulting map showed unambiguous density for the protein including many regions where the search model was locally different from the true structure. The structure could be completed by the application of automated model building methods in phenix.phase_and_build and phenix.autosol (*52*), followed by manual model rebuilding in coot (*53*) in combination with refinement in phenix.refine (*54*) and validation with MolProbity (*55*).

*Structure of bacterial surface layer protein (SLP)*

Excellent diffraction data were available for SLP, but a search for homologues in the PDB using HHpred (*42*) yielded only one hit at a low significance level (E-value of 6.1, sequence identity of 19%) covering only 38% of the protein sequence. Considering that the crystal contains 4 copies of SLP in the asymmetric unit, it was not surprising that molecular replacement attempts failed before the RoseTTAFold models were available.

Initial attempts to solve the structure using an ensemble made from models of the entire protein were partially successful but failed because of crystal packing clashes. However, when the models were divided into two domains, searches with four copies of an ensemble model for the N-terminal domain gave a clear solution with good signal. This turned out to be sufficient to complete the structure if weak phase information from a mercury derivative was added by MR-SAD (*56*). Alternatively, the structure could be solved purely by molecular replacement, by adding four copies of an ensemble model for the C-terminal domain, in which B-factors were computed from the estimated RMS errors and residues with a predicted error greater than 1.3 Å were removed. Automated building procedures were sufficient to complete the structure from this point.

*Structure of secreted fungal protein Lrbp*

Diffraction data were available to 1.53 Å resolution, but no significant hits were found in a search for homologues in the PDB using HHpred (*42*) as the top hit had an E-value of 110. Attempts over the course of 4 years to solve this structure, using a variety of predicted models and small fragments of regular secondary structure had failed.

The initial MR searches using RoseTTAFold models prepared with the default protocol also failed. However, the diversity of the models was increased by varying the selection criteria for the MSA, and the estimated RMS errors were used to delete residues with errors estimated to be greater than 1.3 Å. To generate more diverse models, we collected 8 different MSAs with E-value cutoff of 1e-40, 1e-30, 1e-20, and 1e-10 and sequence coverage cutoff of 50% and 75%. With this strategy, clear solutions for the two copies in the asymmetric unit emerged, leading to a high quality model. As seen in Fig. S3C, the error estimates give a reliable indication of where confidence should be placed in the model.

### **Modeling of GPCR structures**

*GPCR modeling benchmark set construction and evaluation*

A benchmark set of 27 GPCR sequences with experimentally determined structures that were not included in the RoseTTAFold training set was constructed. X-ray and cryo-EM structures determined with resolution higher than 4 Å were excluded. Annotations in the GPCRdb (*8*) were used to classify GPCR sequences, structures, active states, and the transmembrane region residues for analyses. All predicted models were evaluated for the transmembrane regions only. The reference experimental structures were also truncated to the corresponding transmembrane regions, and the TM-score software (*31*) was used to calculate $C_\alpha$ RMSD of the models. To check if templates with similar sequences were available, the sequence identities between the target transmembrane region sequence and the aligned sequences were re-calculated. From the HHblits template search, results with e-value less than 1e-10 were considered, if they were found. The highest sequence identity among the alignments that have transmembrane region coverage higher than 80% was used for analysis. The estimated model accuracy (DAN-lDDT) was predicted by applying the DeepAccNet (*10*) on each truncated model.

*Modeling active and inactive states of GPCRs*

For each target sequence, active and inactive state GPCR template sets were separately provided to two parallel predictions, each generating the corresponding state models. When a template structure in a certain state was not available, models were not predicted for that state. For the benchmark test, templates with sequence identities higher than 70% from HHsearch (*7*) results were excluded to construct the test more fairly.

*GPCR benchmark test performance*

Models with highest estimated accuracy values (DAN-lDDT) were selected for each active and inactive state. RoseTTAFold could predict highly accurate models of both active and inactive states. Examples of good predictions are shown in Fig. S5 panel A and B.

Template-based models of the benchmark set targets were collected from available GPCR model databases. Active state models were brought from GPCRdb (*8*) and inactive state models were downloaded from the Meiler group modeling database (*9*). Targets that could have been modeled easily using any template with sequence identity > 70% in the same state were excluded for analysis. The accuracies of the RoseTTAFold model and corresponding homology model are compared in Fig. S5C. For most of the targets, RoseTTAFold could predict higher TM-score structures.

The best template sequence identity values for each GPCR sequence are reported with estimated model accuracy (DAN-lDDT) and actual accuracy in Fig. S5D. When multiple reference experimental structures existed for the corresponding state, the best $C_\alpha$-RMSD was reported with color representing model accuracy. RoseTTAFold prediction results on the GPCR benchmark set didn't have a high correlation with the best template sequence identity. This again corroborates that the deep-learned network of RoseTTAFold can predict models with accuracies beyond that which can be achieved only with homology information. However, generating highly accurate active state models ($C_\alpha$-RMSD < 1.5 Å) was more feasible when templates with higher sequence identities were available.

The DAN-lDDT of 0.80 can roughly be used as a threshold to discriminate between accurate ($C_\alpha$-RMSD < 1.5 Å or TM-score > 0.9, Fig. S5D) and inaccurate models. Using this guideline to estimate model accuracy could be better applied to inactive state models (Fig. S5D). The active state models turned out to have lower DAN-lDDT than their actual accuracy. The DeepAccNet was trained on monomeric structures only, and the receptor chain in an active state, which would require other chains such as G-proteins as interacting partners, could have been underestimated.

*GPCR models of unknown states*

In the GPCR benchmark set we constructed, 25 targets (as of May 14th, 2021) didn't have known structures of one state, either inactive or active. We predicted models of the unknown state for each target, and models with DAN-lDDT higher than 0.75 were achieved for all targets. These models are provided in http://files.ipd.uw.edu/pub/RoseTTAFold/GPCR_benchmark_one_state_unknown_models.tar.gz

*Human GPCR model generation*

We collected a set of 298 human GPCR sequences without known experimental structures as of May 14th, 2021. Models both in active and inactive states were predicted by applying RoseTTAFold. The best template sequence identity and the estimated accuracy (DAN-lDDT) of the models are reported in Fig. S5D. All models with DAN-lDDT values higher than 0.75 are provided in http://files.ipd.uw.edu/pub/RoseTTAFold/all_human_GPCR_unknown_models.tar.gz. The DAN-lDDT metric can be used to estimate the reliability of each model, and the relative per-residue quality estimation information can be found in the B-factor column.

### Modeling of structurally uncharacterized domains from human proteins

We selected human proteins of biomedical importance based on the number (>50) of literature that are linked to them in Uniprot (*57*) and whether mutations in them are known to cause human diseases according to the DBSAV database (*58*). 7,639 human proteins were selected and domains in them were predicted using HMMER (*59*) search against Pfam database (*60*). A total of 18,233 domains were detected (e-value < 1e-5) in these proteins. Majority of these domains can be modeled confidently by homologous structure in PDB (*61*), and out of the structurally uncharacterized domains, over half of them include a considerably large (> 25%) fraction of residues that are predicted to be disordered (*62*). Excluding domains that are disordered or can be modelled by homology, we removed redundancy, i.e. domains that were mapped to the same Pfam, in the remaining 2,083 domains, resulting in 693 targets to model with our method. We obtained high-quality (estimated lDDT with DeepAccNet (*10*) > 0.8) models for 245 targets (provided in http://files.ipd.uw.edu/pub/RoseTTAFold/human_prot.tar.gz). These models were manually inspected to reveal biological insights with the help of literature, sequence conservation and remote homology that can be detected by searching structurally similar proteins.

For three RoseTTAFold structure models that provided insight into their biological function, their sequences (Q6ICL3:1-259 for TANGO2, P27544:98-304 for CERS1, and Q9BZ11:39-167 for ADAM33 prodomain) were checked against the SWISS-MODEL repository (*61*) for homology models. Their sequences were also submitted to the HHpred server (*63*) for search against the PDB database (PDB_mmCIF70_17_May) and the ECOD (*64*) domain database (ECOD_F70_20200717) using default parameters. For the CERS1 example, where no confident hits were identified, a second MSA generation method using PSI-BLAST against nr70 was used to identify possible template homologs. HHpred results are summarized in Table S3, omitting hits below rank 5. To identify related folds for the examples, RoseTTAFold models were used as queries to search the ECOD database with RUPEE (default settings) (*65*). Potential functional sites for the models were mapped with AL2CO (*66*) using conservations from multiple sequence alignment (MAFFT, default settings (*67*)) of orthologs collected from the OMA database (*68*).

The SWISS-MODEL repository could only generate low quality models for the TANGO2 sequence. However, HHpred generated alignments for several Ntn templates with high confidence (Table S3). We chose the top two templates (3gvz and 2x1d) to generate homology models using the SWISS-MODEL workspace alignment mode (*61*). Each of the homology models was of poor quality based on QMEAN scores (*69*) (-6.12 and -6.11, respectively). These homology models were compared to the RoseTTAFold structure using pairwise DaliLite (*70*) superposition (DaliZ 19.1 and 17.9, respectively). Compared to the RoseTTAFold structure (Fig. S6A), each of the homology models display shifts in alignment and relatively poorly structured loops. Some of the conserved residues that form the RoseTTAFold active site (Fig. S6A, colored red) shifts further away from the active site in each of the homology models: R86, G87, and K166 in the 3gvz model (Fig. S6B) and G49, G51 and K166 in the 2x1d model (Fig. S6C).

Template search for the ADAM33 prodomain confidently identified an incorrect template (4on1_B) corresponding to a fragilysin-3 prodomain fold. While each of the structures possess a similar four-stranded beta-meander, the alpha + beta C-terminus of the fragilysin prodomain extends the beta-meander into a longer sheet (Fig. S14A). Alternately, the N-terminus of ADAM33 continues the beta-meander to form a beta-barrel fold similar to that of lipocalin (Fig. S14B). The HHpred alignment for the fragilysin template incorrectly extended the metalloprotease domain present in both ADAM33 and fragilysin into the prodomain (aligned portions of the prodomains in rainbow, Fig. S14). HHpred search with the ADAM33 prodomain sequence of templates from the ECOD domain database, which separates the prodomain from the metalloprotease, avoids this multi-domain problem.

### Hetero-complex structure prediction using RoseTTAFold

Despite RoseTTAFold being trained on single protein chains, we deployed its ability to make inferences on discontinuous sequence segments to the hetero-oligomer complexes. The only modification we introduced for hetero-complex structure prediction was changes in the positional encoding. We added 200 to the residue numbers of the following subunits to let the network know that it has chain breaks between each subunit.

As a benchmark, we predicted the hetero-oligomer structures of *E. coli.* proteins from the PDB benchmark set (*29*). Among 868 pairs in the PDB benchmark set, we selected 68 interaction pairs having known complex structures of identical or close homologous proteins (sequence identity > 90%) in the PDB and having interface area (calculated by naccess (*71*)) larger than 1,500 Å$^2$. To see whether RoseTTAFold can predict higher order oligomer structures, we also tried to predict hetero-trimer complex structures of bacterial proteins shown in Fig. 4B. For both cases (dimer and trimer prediction), the prediction was made based on a paired alignment of the target complex without any template information.

For human IL12Rβ2-IL12 complex structure prediction, we generated paired alignment by simply pairing the sequences having the same taxonomy ID. Based on this paired sequence alignments and the template structure (IL23R-IL23 complex structure; PDB 6WDQ), the backbone coordinates were predicted using RoseTTAFold. The full-atom structures were generated by FastRelax (*72*) with restraints derived from predicted distances and orientations.

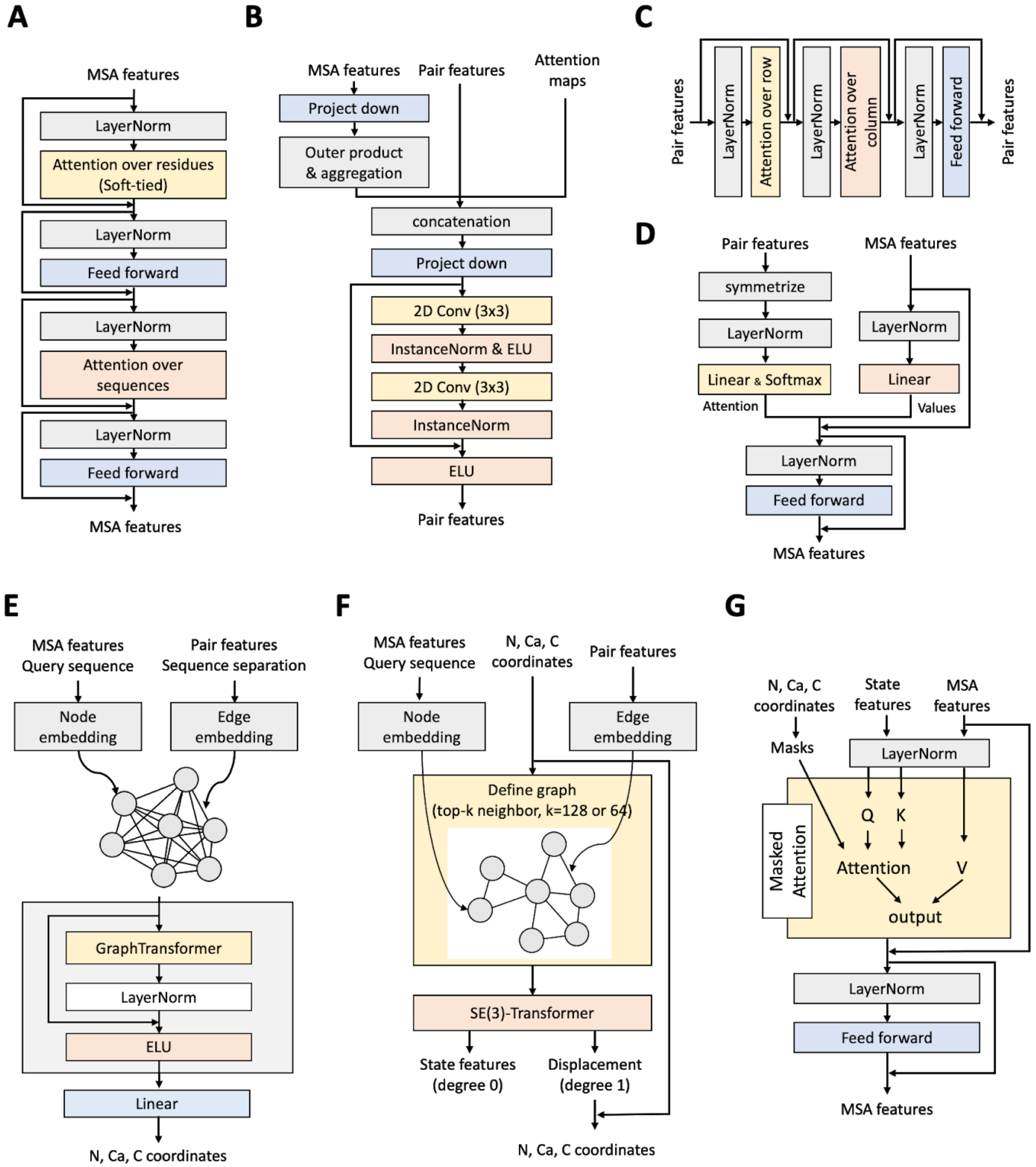


**Fig. S1. Detailed architecture of each component of RoseTTAFold. (A)** MSA updates via self-attention on MSA features. The attention maps over residues are softly tied. **(B)** Pair feature updates based on co-evolution signals derived from MSA features by taking outer-products and weighted averages of them. **(C)** Pair feature refinement through axial attention. **(D)** MSA feature updates based on attention maps derived from given pair features. **(E)** Initial N, $C_\alpha$, C coordinate generation using Graph Transformer architecture. **(F)** 3D coordinate refinements with SE(3)-Transformer. **(G)** MSA feature updates based on given 3D structures using masked attention maps.

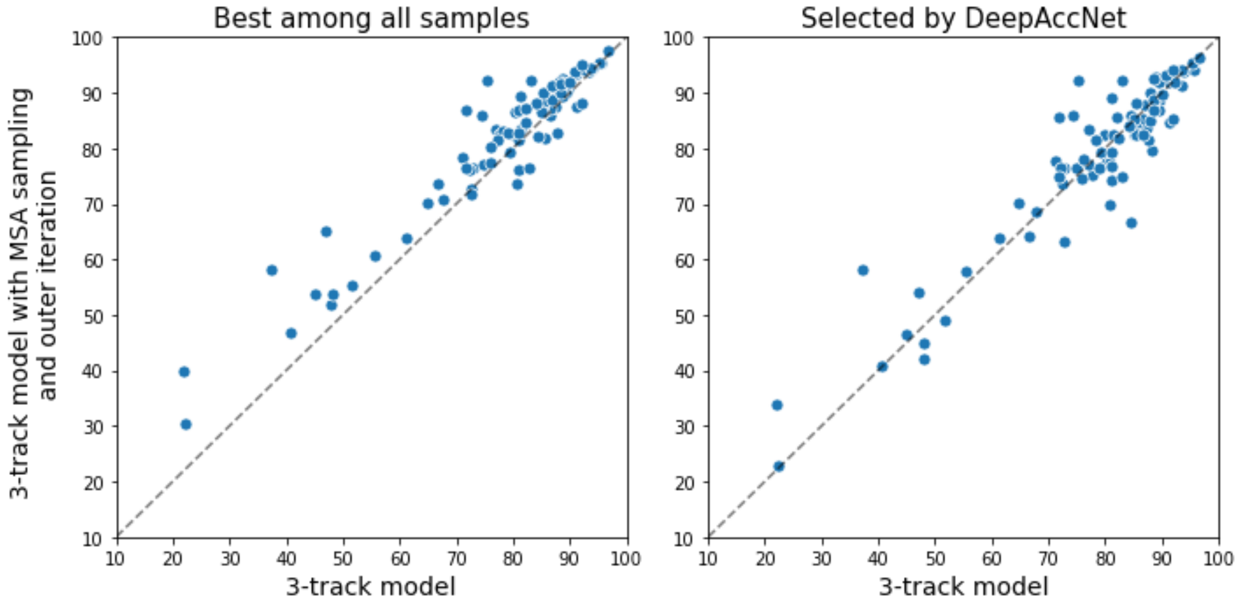


**Fig. S2. Model accuracy changes upon intensive use of the network for inference.** By sampling MSAs randomly and providing predicted structures as templates (y-axis), the 3-track end-to-end model was able to sample much better model structures than the single pass (x-axis) as shown in the left panel. DeepAccNet was able to select improved structures for some cases (right), but there is still room for improvement in model accuracy estimation. The model accuracy is measured by TM-score.

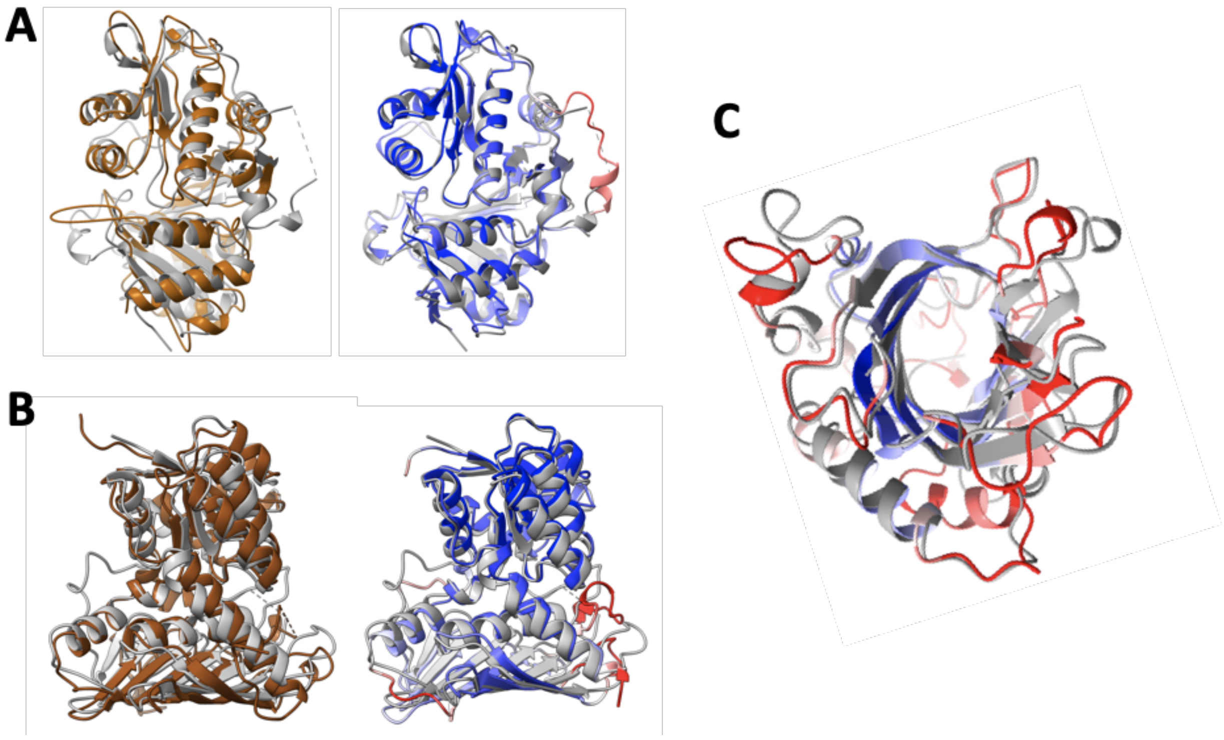


**Fig. S3. Enabling experimental structure determination with RoseTTAFold for proteins having distant homologs. (A)** The final structure of the hexagonal crystal form of GLYAT is shown in gray, with a dashed line representing the disordered loop. The best single template (left) of known structure (PDB entry 1sqh) is shown in brown. The RoseTTAFold model (right) is colored from blue to red by estimated RMS error, ranging from the minimum of 0.56 Å (blue) to 1.5 Å (red). **(B)** Structure determination of an oxidoreductase. The final structure of the oxidoreductase is shown in gray, and the best template (PDB entry 4mkz) is shown in brown. Dashed lines indicate unmodelled residues. The RoseTTAFold model is colored from blue to red by estimated RMS error, ranging from the minimum of 0.6 Å (blue) to 3.5 Å (red). **(C)** The full RoseTTAFold model for Lrbp. The model structure is colored based on estimated RMS error, ranging from the minimum of 0.84 Å (blue) to 1.8 Å (red). The refined structure is shown in gray.

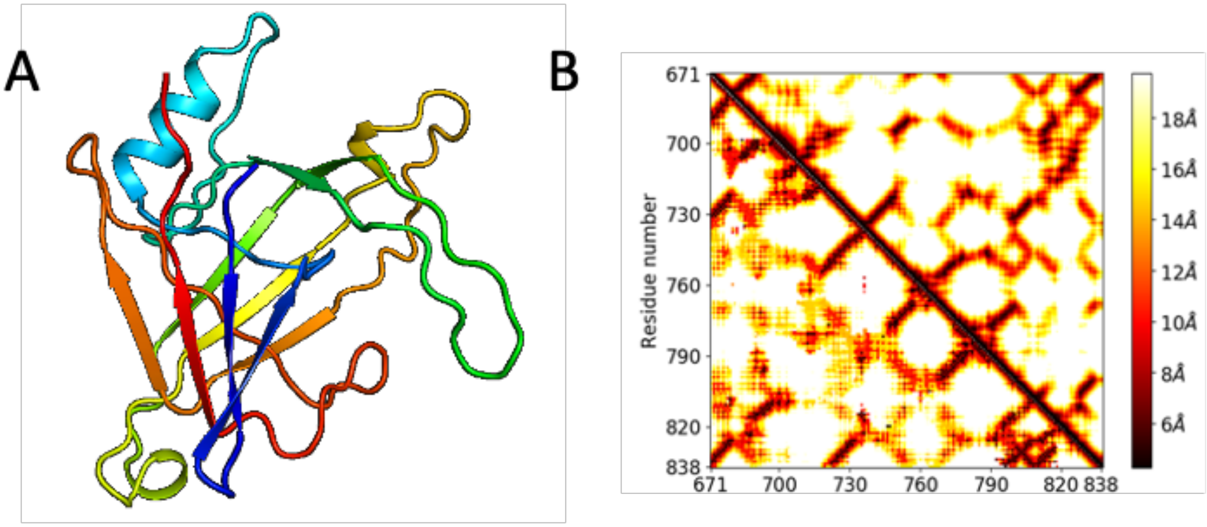


**Fig. S4**. **A trRosetta model for p101 GBD case. (A)** trRosetta predictions led to irregular all-beta topologies that were physically unrealistic and poorly matched to the resulting density. Six-dimensional density map searching did not yield a preferred placement. **(B)** The trRosetta contacts are ambiguous, particularly at longer sequence separations resulting in a totally different fold. The predicted contacts are shown on the lower left triangle and the experimentally determined contacts are on the upper right triangle.

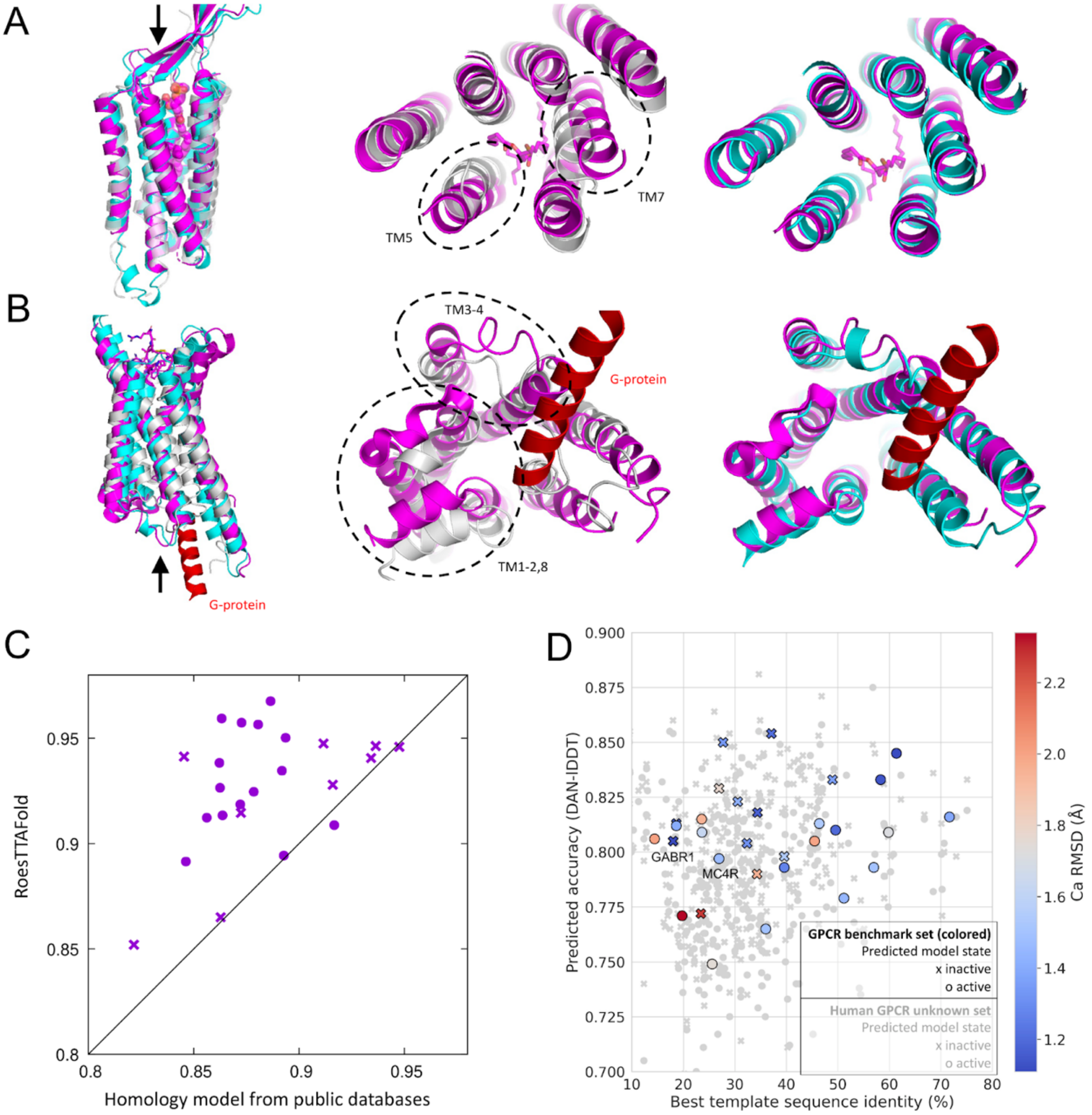


**Fig. S5. GPCR modeling. (A, B)** Models built for GPCRs not in the training set are compared to crystal structures. **(A)** The best DAN-lDDT (0.81) inactive state model of GABR1_HUMAN (cyan, Uniprot ID Q9UBS5) compared to the native (PDB 6w2y chain B, magenta) and the closest homolog of known structure (PDB 4or2 chain A, gray, seqID 18%). Transmembrane region $C_\alpha$-RMSD was 1.14 Å. Middle and right panels focused on extracellular regions (top view). **(B)** The best DAN-lDDT (0.80) active state model of MC4R_HUMAN (cyan, Uniprot ID P32245) compared to the native (PDB 7aue chain R, magenta, G-protein helix in red) and the closest homolog of known structure (gray, PDB 3kj6 chain A, seqID 27%). Transmembrane region $C_\alpha$-RMSD was 1.49 Å. Middle and right panels focused on intracellular regions (bottom view). **(C)** Accuracies (in TM-score) of RoseTTAFold models versus template-based models from public databases (*8*, *9*). Only transmembrane regions were considered. **(D)** For each active (o) and inactive (x) state prediction, the best template sequence identity and predicted model accuracy (DAN-lDDT) are reported. The color gradient represents actual model accuracy in $C_\alpha$-RMSD for the subset of proteins of known structure, ranging from 1.2 Å (accurate, blue) to 2.2 Å (inaccurate, red). The human GPCR set with unknown structures is shown in light gray. Data with DAN-lDDT between 0.7 and 0.9 are only shown.

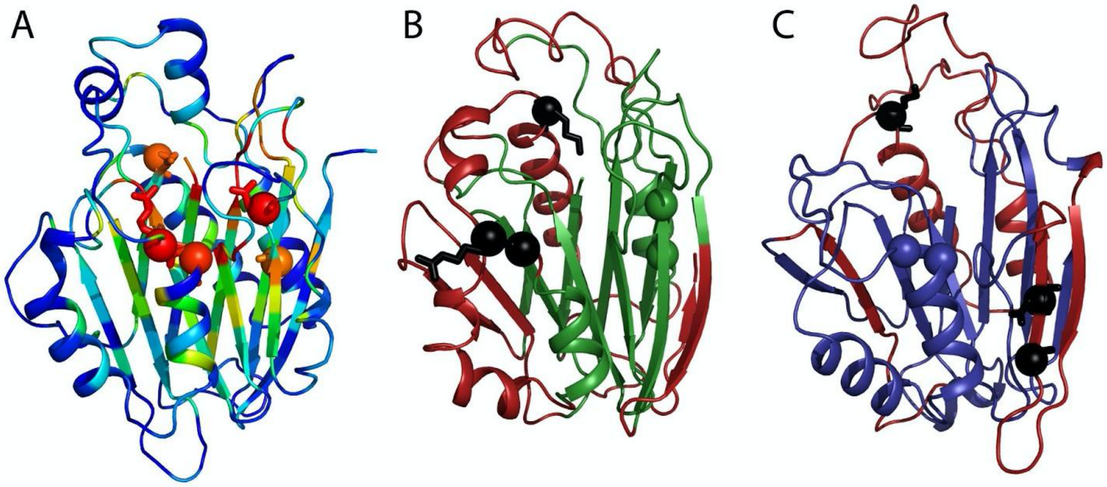


**Fig. S6. RoseTTAFold structure for TANGO2 improves homology models. (A)** TANGO2 RoseTTAFold structure is colored by ortholog conservation in the rainbow from variable (blue) to conserved (red). Shifted active site residues in either of the homology models are shown in stick with the $C_\alpha$ in sphere. **(B)** Homology model based on the top HHpred hit to 3gvz template is colored green (aligned with the RoseTTAFold structure) or red (shifted alignment). Three conserved residues (black sphere and stick) shift away from the active site. **(C)** Homology model based next best HHpred hit to 2x1d template is colored blue (aligned with the RoseTTAFold structure) or red (shifted alignment). Three conserved residues (black sphere and stick) shift away from the active site.

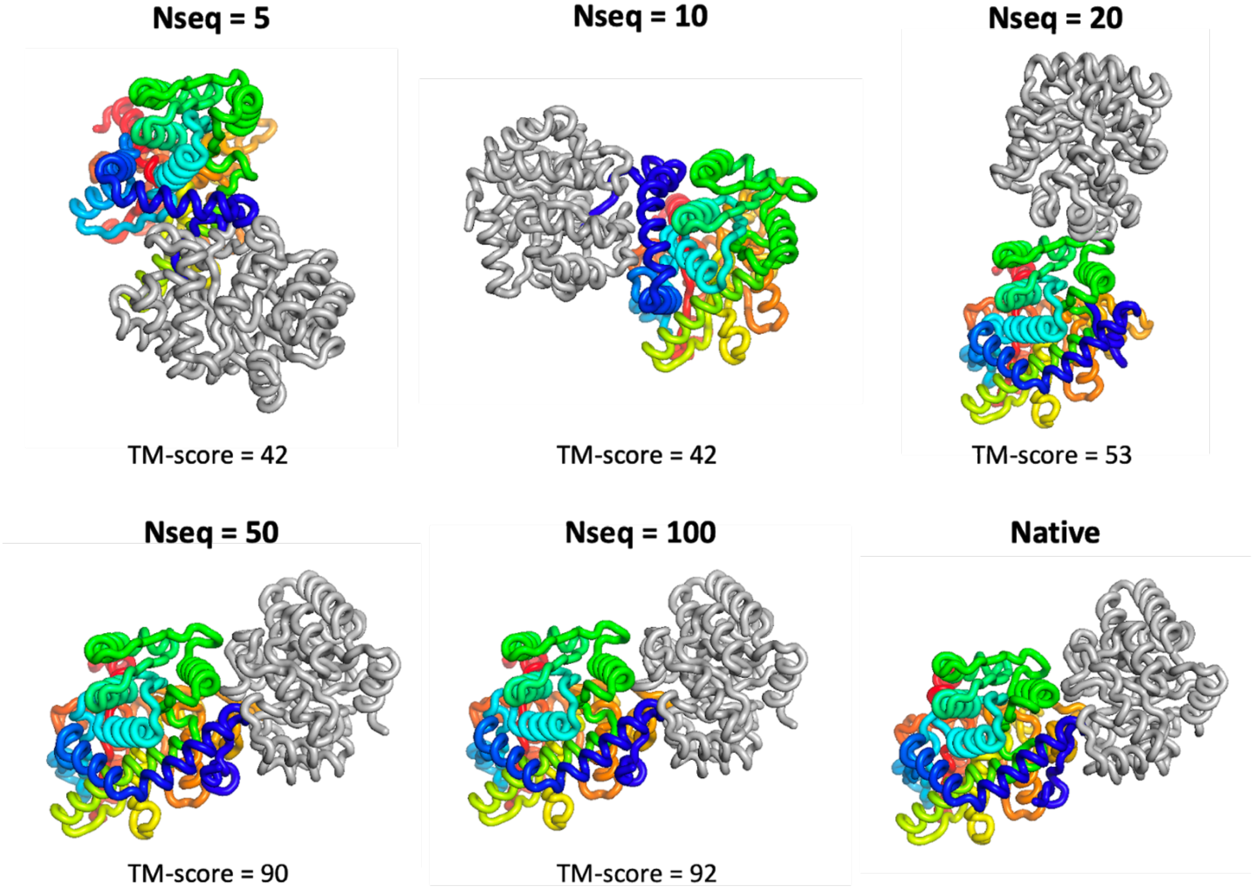


**Fig. S7**. **Complex model accuracy depends on the number of sequences in paired MSA**. The number of sequences in the MSA is written on the top while complex TM-score of the predicted model is written on the bottom.

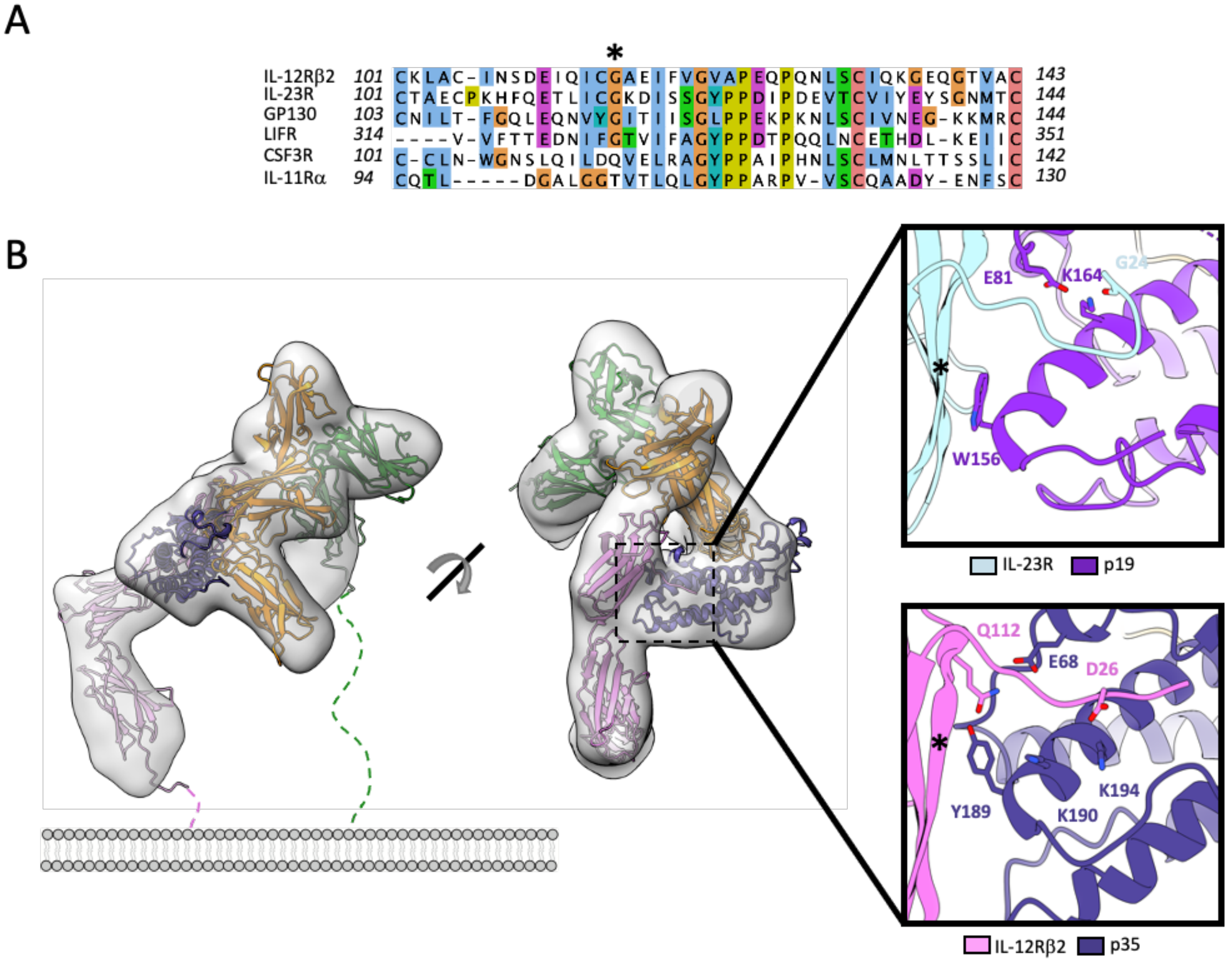


**Fig. S8. Analysis of the complete IL-12 receptor complex model. (A)** Multiple sequence alignment of gp130 family cytokines highlighting the conserved glycine residue in IL-12Rb2 (115), IL-23R (116), GP130 (117), and LIFR (324). Residues were colored using ClustalX. **(B)** Experimental cryoEMdensity of the quaternary IL-12 complex (EMD-21645) fit with IL-12Rb2 model. Inset shows a comparison of the interaction between IL-23R and p19 (top, PDB: 6WDQ) and IL-12Rb2 and p35 (bottom, computational model). Star represents position of glycine residue (G115 in IL-12Rb2, G116 in IL-23R).

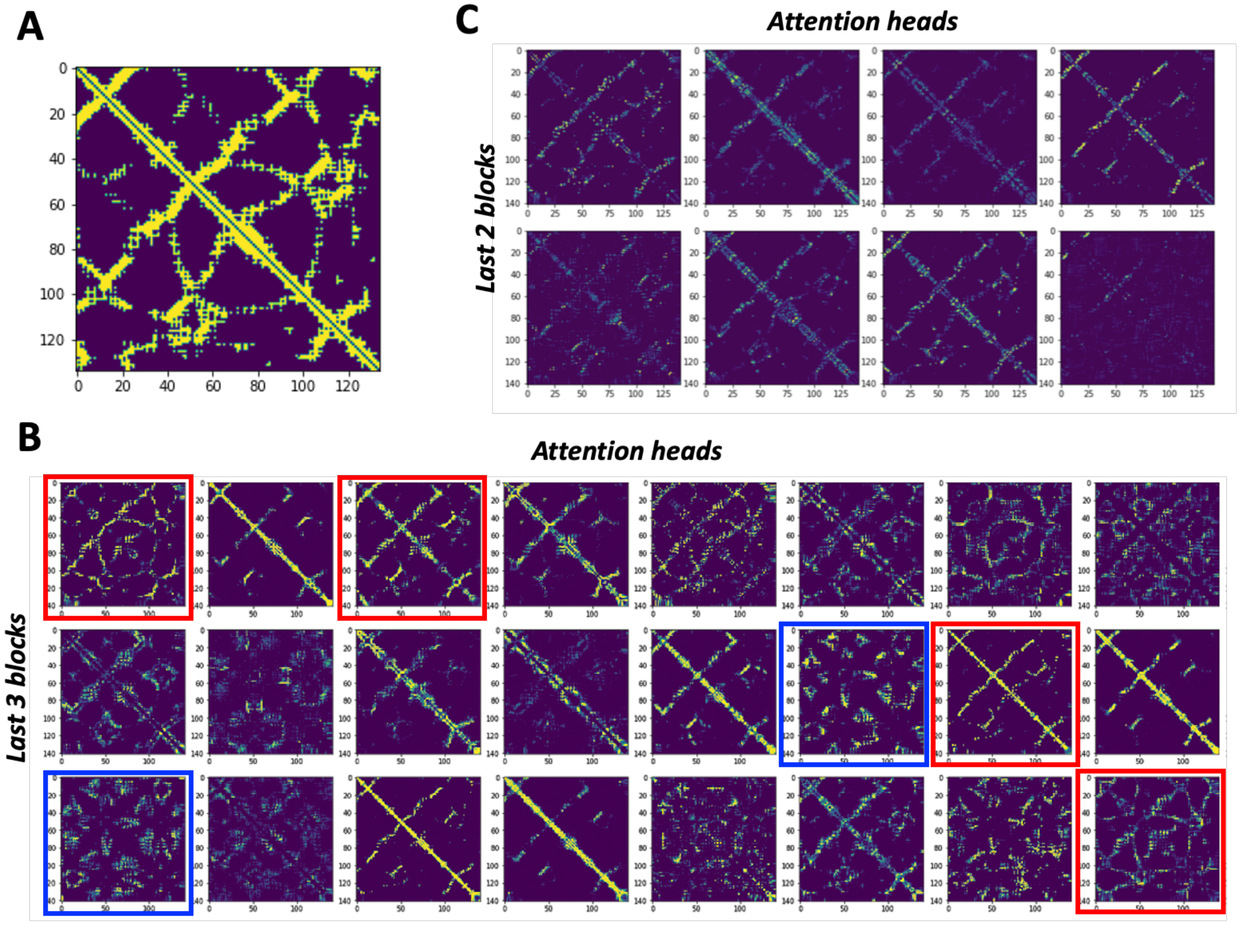


**Fig. S9. Examples of attention maps used to update MSA. (A)** True contact map of CASP14 target T1049. **(B)** Attention maps from self-attention on MSA features for the last three blocks of the 2-track model (76M parameter model). Some of the attention heads (red boxes) resemble a true contact map. For some cases (blue boxes), it only attends to the positions not making the direct contacts. **(C)** Attention maps derived from pair features used to update MSA features. It also shows the similar pattern to the true contact map. The attention maps shown in this figure are symmetrized for clear visualization.

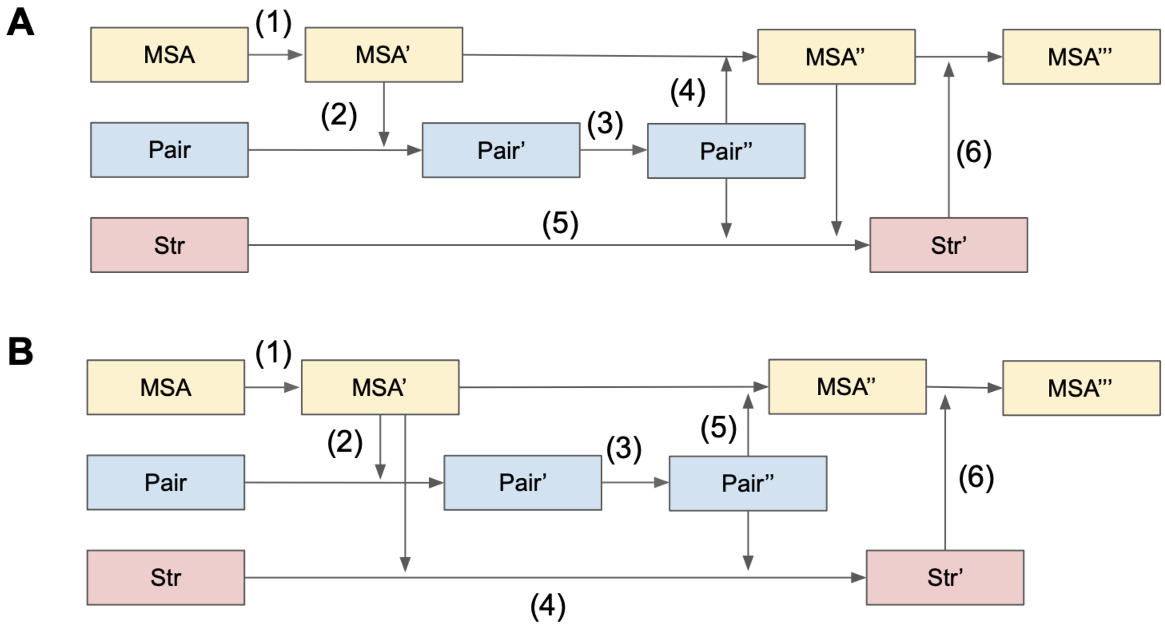


**Fig. S10. Two different 3-track block definitions. (A)** MSA and pair features are synchronized before structure updates. **(B)** Structure is updated based on unsynchronized MSA and pair features. The numbers in parentheses indicate the order of calculation.

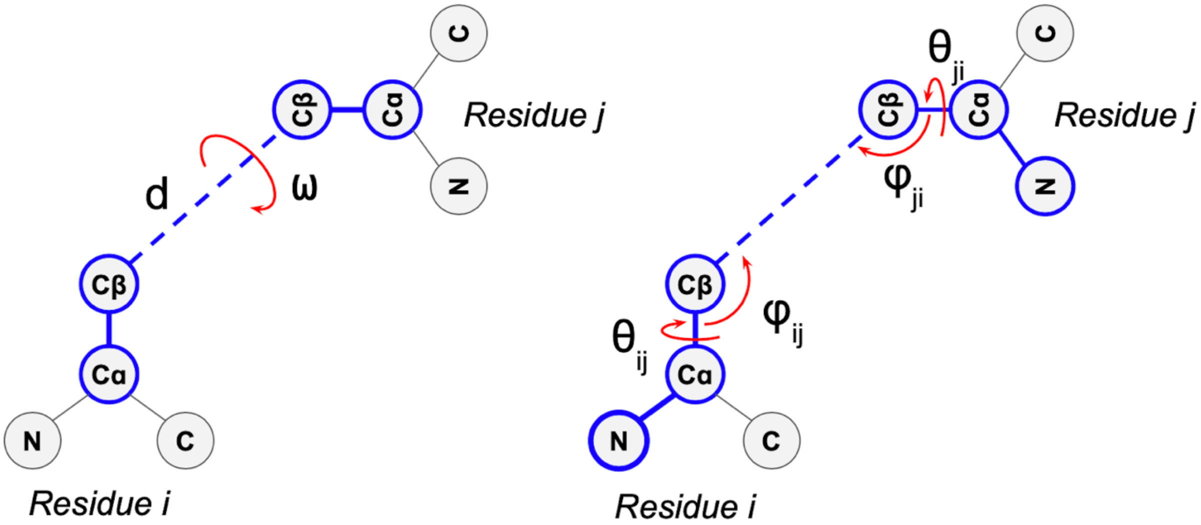


**Fig. S11. 6D representation of rigid body transform between two residues.** It includes distance (d) between Cβ atoms, dihedral angle (w) arong virtual bond connecting two Cβ atoms, and two dihedral angles ($\theta_{ij}$, $\theta_{ji}$) and two pseudo-bond angles ($\varphi_{ij}$, $\varphi_{ji}$) specifying the direction of the Cβ atom of a residue in a reference frame centered on the other residue.

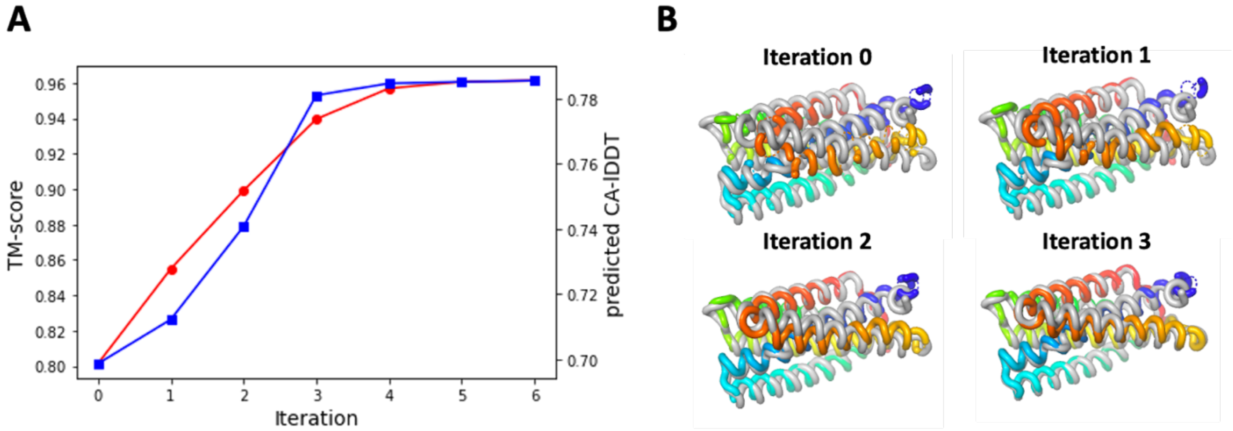


**Fig. S12. An example (T1024-D1 from CASP14 targets) of Iterative refinement using SE(3)-Transformers. (A)** Model accuracy (TM-score) is improved with iterative refinement. Predicted $C_\alpha$-lDDT from the network shows a good correlation to the actual model accuracy. **(B)** The model structure at each iteration is shown. The RoseTTAFold models are colored in rainbow (blue; N-terminal, red; C-terminal) while the native structures are colored in gray.

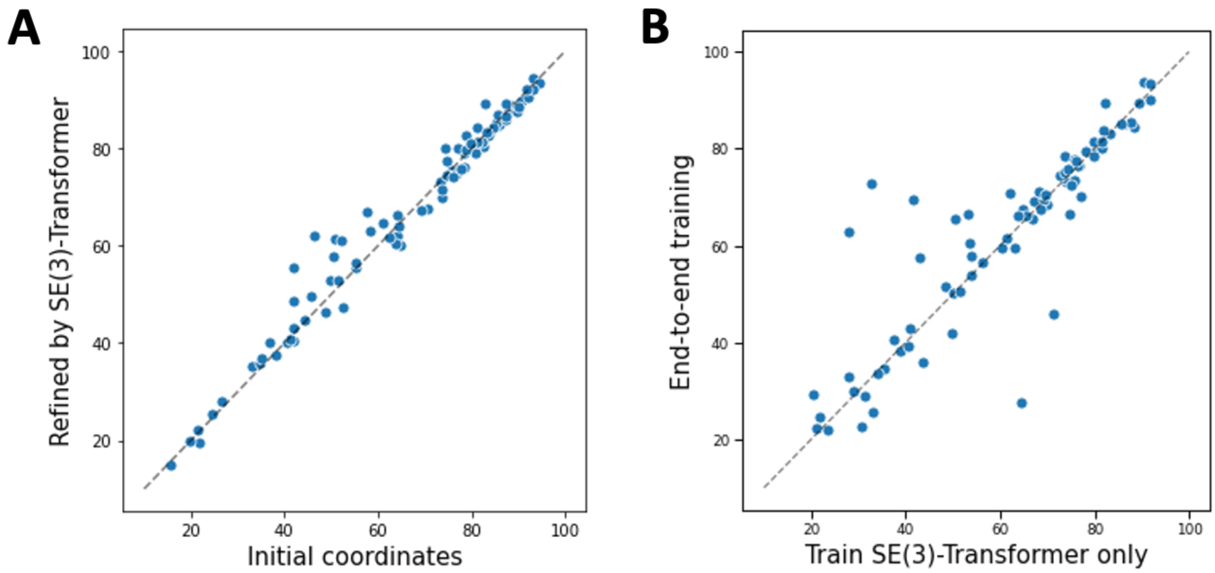


**Fig. S13. Experiments with SE(3)-Transformer layers on top of the two track model. (A)** Model accuracy comparison between initial coordinates generated by the simple graph-based network and the refined models through SE(3)-Transformer. **(B)** Model accuracy comparison between networks trained in two different ways: SE(3)-Transformer trained separately with the frozen 2-track model (x-axis) and structure module having the same architecture trained together with 2-track model part (y-axis). Model accuracy is measured in terms of TM-score.

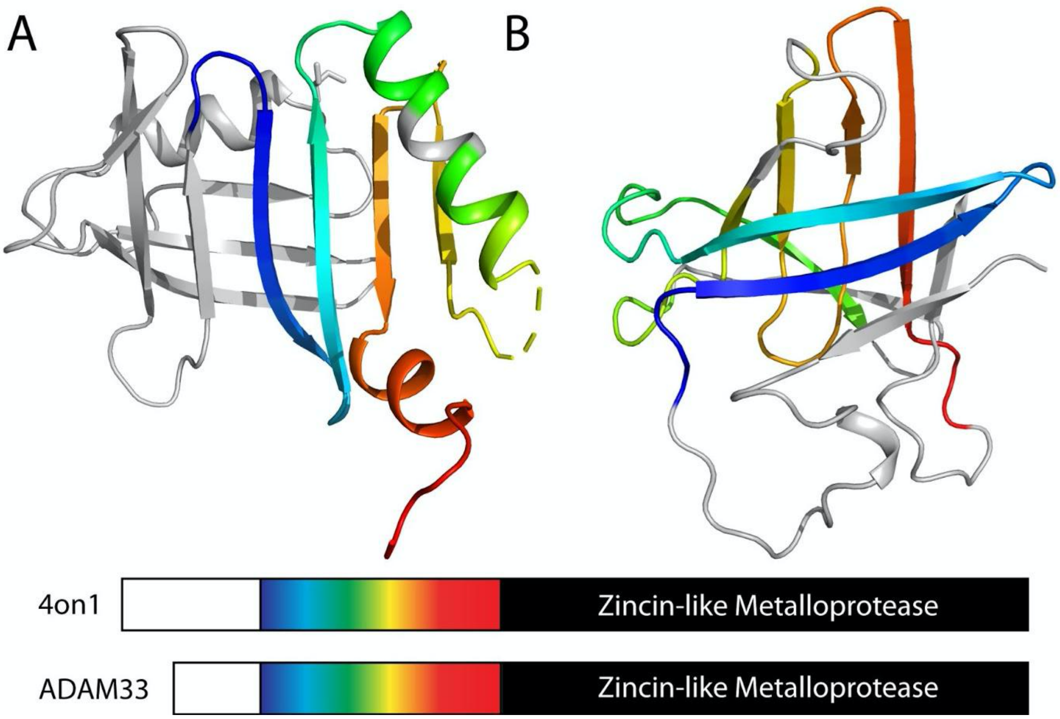


**Fig. S14. RoseTTAFold structure avoids multi-domain problems. (A)** The prodomain from HHpred template 4on1 is in ribbon and adopts a fragilysin-like alpha + beta fold with a central 4-stranded beta-meander. The domain architecture below highlights a C-terminal metalloprotease that is also in ADAM33. The HHpred template alignment incorrectly extends into the prodomain (aligned sequence in rainbow). **(B)** ADAM33 RoseTTAFold structure (oriented by its corresponding central beta-meander) adopts a lipocalin-like beta-barrel. The aligned beta-meander sequence (in rainbow) is unrelated to the alpha + beta sequence from the template.

**Table S1**. Performance of different model architectures in terms of inter-residue geometry prediction loss (cross entropy), top L long-range contact accuracy and $C_\alpha$-lDDT.

| Architecture | Inter-residue geometry loss | Top L long-range contact accuracy | $C_\alpha$-lDDT |
|---|---|---|---|
| Single Track (Sequential processing of MSA and pair feature) | | | |
| Architecture 1) Hand-crafted features + 2D convolution | 5.56 | 54% | - |
| Architecture 2) MSA encoder + 2D convolution | 5.49 | 56% | - |
| Architecture 3) MSA encoder + Axial attention | 5.14 | 58% | - |
| 2-track (Parallel track for MSA and pair features) | | | |
| Architecture 4) Untied + addition + cross | 5.54 | 54% | - |
| Architecture 5) Untied + addition + direct | 5.18 | 58% | - |
| Architecture 6) Untied + concat + direct | 5.01 | 60% | - |
| Architecture 7) Soft-tied + concat + direct | 4.84 | 62% | - |
| Architecture 8) architecture 7 + scale-up | 4.50 | 67% | - |
| Architecture 9) architecture 8 + SE(3) structure module | 4.54 | 67% | 0.70 |
| 3-track (Parallel track for MSA, pair, and 3D coordinates) | | | |
| Architecture 10) Structure update w/ unsynchronized MSA and pair features (**Fig S10B)** | 4.63 | 64% | 0.68 |
| Architecture 11) Structure update w/ synchronized MSA and pair features (**Fig S10A)** | 4.36 | 69% | 0.72 |
| Architecture 12) architecture 11 + SE(3) structure module | 4.39 | 69% | 0.77 |

**Table S2**. Current refinement statistics for crystal structures

| Crystal | GLYAT | Oxidoreductase | SLP | Lrbp |
|---|---|---|---|---|
| Space group | $P6_5$ | $P2_1$ | $P2_12_12_1$ | $P2_1$ |
| Cell dimensions | | | | |
| *a, b, c* (Å) | 97.18, 97.18, 144.63 | 79.15, 157.86, 95.01 | 63.16, 98.87, 155.12 | 50.10, 81.37, 78.47 |
| *α, β, γ* (˚) | 90, 90, 120 | 90, 114.45, 90 | 90, 90, 90 | 90, 107.57, 90 |
| Resolution (Å) | 1.65 | 2.34 | 2.18 | 1.53 |
| No. non-H atoms | 5002 | 14568 | 5463 | 4621 |
| No. reflections | 83145 | 87002 | 49958 | 89331 |
| $R_{work}$, $R_{free}$ | 0.174, 0.200 | 0.283, 0.322 | 0.216, 0.250 | 0.248, 0.280 |

**Table S3**. HHpred Results Summary for TANGO2, CERS1, and ADAM33

| Example | Sequence | MSA | Hit | Prob | Cols | Query | Temp. | Coverage |
|---|---|---|---|---|---|---|---|---|
| TANGO2 | Q6ICL3:1-259 | Uniref30 | PDB: 3GVZ_A | 98.9 | 219 | 1-252 | 25-256 | 0.846 |
| TANGO2 | Q6ICL3:1-259 | Uniref30 | ECOD: e3gvzA1 | 98.5 | 217 | 2-253 | 1-232 | 0.838 |
| TANGO2 | Q6ICL3:1-259 | Uniref30 | PDB: 2X1D_D | 98.2 | 210 | 1-254 | 102-330 | 0.811 |
| TANGO2 | Q6ICL3:1-259 | Uniref30 | PDB: 3HBC_A | 97.8 | 217 | 1-255 | 3-274 | 0.838 |
| TANGO2 | Q6ICL3:1-259 | Uniref30 | ECOD: e3hbcA1 | 97.7 | 212 | 1-247 | 3-268 | 0.819 |
| CERS1 | P27544:98-304 | Uniref30 | ECOD: e3nqwB1 | 8.5 | 53 | 64-116 | 12-65 | 0.256 |
| CERS1 | P27544:98-304 | PDB70 | PDB: 6TY2_A | 17.7 | 22 | 103-124 | 27-48 | 0.106 |
| ADAM33 | Q9BZ11:39-167 | Uniref30 | PDB: 4ON1_B | 96.3 | 90 | 24-117 | 89-184 | 0.698 |
